# Helper NLRs Nrc2 and Nrc3 act co-dependently with Prf/Pto and activate MAPK signaling to induce immunity in tomato

**DOI:** 10.1101/2023.08.17.553711

**Authors:** Ning Zhang, Joyce Gan, Lauren Carneal, Juliana González-Tobón, Melanie Filiatrault, Gregory B. Martin

## Abstract

Plant intracellular immune receptors, primarily nucleotide-binding, leucine-rich repeat proteins (NLRs), can detect virulence proteins (effectors) from pathogens and activate NLR-triggered immunity (NTI). Recently, ‘sensor’ NLRs have been reported to function with ‘helper’ NLRs to activate immune responses. We investigated the role of two helper NLRs, Nrc2 (NLR required for cell death 2) and Nrc3, on immunity in tomato to the bacterial pathogen *Pseudomonas syringae* pv. *tomato* (*Pst*) mediated by the sensor NLR Prf and the Pto kinase. Loss-of-function mutations in both *Nrc2* and *Nrc3* completely compromised Prf/Pto-mediated NTI to *Pst* containing the cognate effectors AvrPto and AvrPtoB. An *nrc3* mutant showed intermediate susceptibility between wild-type tomato plants and a *Prf* mutant, while an *nrc2* mutant developed only mild disease symptoms. These observations indicate that Nrc2 and Nrc3 act additively to contribute to Prf/Pto-mediated immunity. We also examined at what point Nrc2 and Nrc3 act in the Prf/Pto-mediated immune response. In the *nrc2/3* mutant, programmed cell death (PCD) normally induced by constitutively-active variants of AvrPtoB, Pto or Prf was abolished, but that induced by M3Kα or Mkk2 was not. PCD induced by a constitutively active variant of Nrc3 was also abolished in a *Nicotiana benthamiana* line with reduced expression of *Prf*. MAPK activation triggered by expression of AvrPto in the wild-type Pto-expressing tomato plants was completely abolished in the *nrc2*/3 mutant. These results indicate that Nrc2 and Nrc3 act in concert with Prf/Pto and upstream of MAPK signaling. Nrc2 and Nrc3 were not required for the HR triggered by Ptr1, another sensor NLR mediating *Pst* resistance, although these helper NLRs do appear to be involved in resistance to certain *Pst* race 1 strains.

## Introduction

Plants have evolved a complex, two-layered immune system to defend against themselves pathogens. Upon attempted infection, plants use extracellular pattern recognition receptors (PRRs) to detect the presence of conserved microbe-associated molecular patterns (MAMPs), such as bacterial flagellin-derived peptides, initiating PRR-triggered immunity (PTI) (Dangl et al., 2013; Saijo et al., 2018; DeFalco and Zipfel, 2021). Plants also have a second robust immune response, driven by intracellular immune receptors, primarily nucleotide-binding, leucine-rich repeat proteins (NLRs). NLRs detect pathogen virulence proteins (effectors) and activate NLR-triggered immunity (NTI), also known as effector-triggered immunity (ETI) (Qi and Innes, 2013; Cui et al., 2015; Lolle et al., 2020; Kourelis and Adachi, 2022).

Activation of NLR proteins in plants is typically initiated when an effector protein is recognized either directly by the leucine-rich repeat (LRR) domain of the NLR or indirectly via the modification of plant proteins that the NLR monitors, mechanisms described by the ‘guard’ and ‘decoy’ models (Tang et al., 1996; Zhou et al., 1998; Belkhadir et al., 2004; Shabab et al., 2008; van der Hoorn and Kamoun, 2008; Wu et al., 2015). Recent studies revealed that upon activation, at least some NLR proteins undergo oligomerization, leading to the formation of larger complexes known as resistosomes (Wang et al., 2019; Martin et al., 2020; Wang et al., 2020; Forderer et al., 2022; Kourelis and Adachi, 2022; Kourelis et al., 2022; Zhao et al., 2022; Ahn et al., 2023a; Contreras et al., 2023). The resistosomes trigger downstream responses, including but not limited to the production of reactive oxygen species (ROS), transcriptional reprogramming, and eventually programmed cell death (PCD), also known as the hypersensitive response (HR), which restricts pathogen growth at the site of infection (Pombo et al., 2014; Balint-Kurti, 2019). Collectively these responses confer disease resistance. It remains unknown whether all NLRs, particularly those with less conserved protein motifs, form resistosomes upon activation (Lolle et al., 2020; Wang et al., 2020; Forderer et al., 2022; Kourelis and Adachi, 2022)

Plants harbor a large and diverse repertoire of NLRs, ranging from 50 to 1000 genes per species (Shao et al., 2016). NLRs are broadly categorized into three subclasses based on their variable N-terminal domains: TIR-NLRs (TNL proteins) with the toll and interleukin-1 receptor (TIR) domain, CC-NLRs (CNL proteins) with the Rx-type coiled-coil (CC) domain, and RPW8-NLRs (RNL proteins) with the RPW8-type CC (CC_R_) domain (Collier et al., 2011). Based on their functionality, plant NLRs can also be grouped into ‘sensor’ NLRs that directly or indirectly detect pathogen effectors and ‘helper’ NLRs that are not directly involved in effector recognition but rather may mediate downstream immune signaling from multiple sensor NLRs (Derevnina et al., 2021; Adachi et al., 2023). Emerging studies indicate that some plant NLRs function in pairs or networks, rather than as single units responsible for both recognizing effectors and activating subsequent downstream responses (Wu et al., 2017; Adachi et al., 2019; Dillon et al., 2019; Kourelis and Adachi, 2022; Shimizu et al., 2022). This model suggests a more complex system of plant-microbe interactions where multiple NLR proteins work together to coordinate pathogen detection and signaling, enhancing the versatility and specificity of plant immune responses (Adachi et al., 2019). One helper NLR protein family, known as <u>N</u>LR <u>r</u>equired for <u>c</u>ell death (*Nrc*) (Gabriels et al., 2007), present in *Solanaceous* plants such as tomato (*Solanum lycopersicum*), *Nicotiana benthamiana*, potato (*Solanum tuberosum*), pepper (*Capsicum annuum*), and eggplant (*Solanum melongena*), are required by many CNL or TNL proteins to defend against diverse pathogens including bacteria, viruses, oomycetes, nematodes, and insects (Wu et al., 2016; Wu et al., 2017; Adachi et al., 2019; Wu and Kamoun, 2021; Kourelis et al., 2022; Shimizu et al., 2022; Adachi et al., 2023; Ahn et al., 2023a).

The interaction of tomato (*Solanum lycopersicum*) with the bacterial pathogen *Pseudomonas syringae* pv. *tomato* (*Pst*) is a model system for understanding the molecular basis of bacterial pathogenesis and plant immunity (Pedley and Martin, 2003; Oh and Martin, 2011a; Martin, 2012). The tomato CNL protein Prf has been particularly well characterized for its role in tomato immunity to bacterial speck disease (Salmeron et al., 1996; Rathjen et al., 1999; Pedley and Martin, 2003; Mucyn et al., 2006; Balmuth and Rathjen, 2007; Gutierrez et al., 2010; Kud et al., 2013; Ntoukakis et al., 2013; Saur et al., 2015; Sheikh et al., 2023). Prf forms an immune complex with an intracellular serine/threonine kinase protein Pto, which can recognize and bind to the effectors AvrPto and AvrPtoB from race 0 strains of *Pst* such as DC3000 (Balmuth and Rathjen, 2007; Ntoukakis et al., 2013; Saur et al., 2015). Recognition of AvrPto/AvrPtoB by the Prf/Pto complex triggers NTI responses including activation of MAPK signaling cascades and eventually HR-associated cell death (Tena et al., 2001; Ekengren et al., 2003; del Pozo et al., 2004; Pedley and Martin, 2004; Oh et al., 2010; Oh and Martin, 2011b). In tomato, two *MAPKKK* genes (*M3Kα* and *M3Kε*), two *MAPKK* genes (*Mkk1* and *Mkk2*), and two *MAPK* genes (*Mpk2* and *Mpk3*) play important roles in Prf/Pto-mediated immune signaling against *Pst* infection (Ekengren et al., 2003; del Pozo et al., 2004; Pedley and Martin, 2004; Melech-Bonfil and Sessa, 2010). The tomato *M3Kα*, a member of subgroup A2 in the MEKK family, was initially identified by a virus-induced gene silencing (VIGS) screen in *N. benthamiana* (del Pozo et al., 2004). Expression of M3Kα in leaves of *N. benthamiana* activates Mpk2 and Mpk3 and results in PCD (del Pozo et al., 2004; Pedley and Martin, 2004). A screen for M3Kα-interacting proteins in tomato identified a 14-3-3 protein, Tft7, which positively regulates Prf/Pto-mediated PCD in tomato and also interacts with Mkk2, which acts downstream of M3Kα (Oh et al., 2010; Oh and Martin, 2011b). Recently, two other 14-3-3 proteins, Tft1 and Tft3, were identified as interacting partners of the M3Kα protein and the Prf/Pto complex (Sheikh et al., 2023). Another tomato M3Kα-interacting protein, Mai1, is a receptor-like cytoplasmic kinase, that acts upstream of M3Kα and positively contributes to PCD and disease resistance associated with Prf/Pto activation (Roberts et al., 2019).

Tomato has nine *Nrc* helper proteins, with seven (Nrc1, Nrc2, Nrc3, Nrc4a, Nrc4b, Nrc4c, and NrcX) mediating NTI responses along with multiple sensor NLR proteins (Sueldo et al., 2015; Wu et al., 2016; Wu et al., 2017; Wu et al., 2020; Wu and Kamoun, 2021; Kourelis et al., 2022; Adachi et al., 2023; Sheikh et al., 2023). For example, reduction of *Nrc2* and *Nrc3* gene expression by VIGS in *N. benthamiana* and tomato significantly compromised Prf/Pto-mediated resistance to *Pst* expressing AvrPto and AvrPtoB (Wu et al., 2016; Wu and Kamoun, 2021). Recently there has been significant progress in elucidating the mechanism underlying the interaction between these helper NLRs and the Prf/Pto complex during the NTI response (Sheikh et al., 2023). Nrc2 and Nrc3 were found to directly interact with Prf in unchallenged leaves. Upon effector recognition, Nrc2 and Nrc3 dissociate from the Prf/Pto complex, triggering downstream responses that ultimately lead to programmed cell death and the restriction of pathogen growth (Sheikh et al., 2023). Here, to further investigate the role of Nrc2 and Nrc3 in Prf/Pto-mediated immunity, we used CRISPR/Cas9 to generate tomato lines with loss-of-function mutations in *Nrc2* and *Nrc3.* Single nrc2 and nrc3 mutants, and double nrc2/3 mutants were then tested for their response to *Pst* strains expressing either one or both of the effectors AvrPto and AvrPtoB and used to investigate at what point Nrc2 and Nrc3 act in the Prf/Pto-mediated immune signaling pathway. Additionally, we used a *N. benthamiana* line with knocked-down expression of *Prf* to assess the requirement of Prf for Nrc3-mediated cell death.

## Results

### Generation of loss-of-function mutations in five *Nr*c genes in tomato

In unchallenged wild-type tomato plants, the five *Nrc* genes *Nrc1*, *Nrc2*, *Nrc3*, *Nrc4a* and *Nrc4c,* exhibit distinct expression patterns (Supplemental Fig. S1). During the *Pst*-associated PTI response in tomato leaves the transcript abundance of each gene is significantly increased, whereas the transcript abundance of just *Nrc2* and *Nrc4a* are increased during the *Pst*-associated NTI response (Supplemental Table S1). The basal transcript abundance of *Nrc3* is the highest among these five *Nrc* genes (Supplemental Table S1). To better understand the role of the *Nrc* genes in plant immunity, we used CRISPR/Cas9 to generate mutations in *Nrc1*, *Nrc2*, *Nrc3*, and *Nrc4a* and *Nrc4c* (Supplemental Fig. S2). After transformation of the tomato cultivar Rio-Grande PtoR (RG-PtoR, which has the *Prf* and *Pto* genes), we obtained multiple T0 mutants disrupting each target gene, from which two independent homozygous mutants of each gene were derived and used in this study. Insertions or deletions in each of the *Nrc* mutant alleles, except for the nrc3-2 allele, resulted in a predicted premature stop codon near the N-terminus of each protein and thus they are likely loss-of-function mutations. As Nrc2 and Nrc3 were previously reported to function redundantly (Wu et al., 2016; Wu and Kamoun, 2021), we crossed the nrc2-1 and nrc2-2 mutants with the nrc3-1 mutant and generated two nrc2/3 double mutant lines (Supplemental Fig. S2). The growth, development, and overall morphology of each nrc mutant plant was indistinguishable from wild-type RG-PtoR plants although there appeared to be a slight effect of the combined nrc2/3 mutations on fruit weight and width (Supplemental Fig. S3).

### Nrc2 and Nrc3 act redundantly and also additively in Prf/Pto-mediated immunity

To test whether the mutation in any of *Nrc* genes affects the tomato response to *Pseudomonas syringae* pv. *tomato* (*Pst*), we vacuum infiltrated each of the nrc mutants, wild-type RG-PtoR, and RG-prf3 plants (containing a mutation in *Prf* that makes the Prf/Pto pathway nonfunctional) with the *Pst* strain DC3000 (1×10^6^ cfu/mL) that expresses AvrPto and AvrPtoB which together activate a strong Prf/Pto-mediated NTI (Fig. 1 and Supplemental Fig. S4; (Kvitko et al., 2009; Lin and Martin, 2005)). To simplify, we show in the main Figures results for only the mutants nrc1-1, nrc2-1, nrc3-1, nrc4a/c-1, and nrc2/3-1 (hereafter referred to as nrc1, nrc2, nrc3, nrc4a/c, nrc2/3) and results of the other mutants (nrc1-2, nrc2-2, nrc3-2, nrc4a/c-2, and nrc2/3-2) in the Supplemental section. Three days after inoculation with DC3000, there were no distinguishable differences in disease symptoms or bacterial populations between wild-type RG-PtoR plants and the nrc1, nrc2, nrc3, or nrc4a/c mutants (Fig. 1). However, the nrc2/3 double mutant, similar to the susceptible RG-prf3 line, displayed a complete loss of resistance to DC3000. Bacterial populations in nrc2/3 were similar to those in RG-prf3 and were 10-fold more than in other nrc mutants and RG-PtoR plants. Comparable disease symptoms and bacterial populations were observed in the other independent mutant lines (Supplemental Fig. S4). These findings suggest that Nrc2 and Nrc3 contribute redundantly to Prf/Pto-mediated NTI, whereas Nrc1 and Nrc4a/c do not appear to contribute to this response as reported earlier in *N. benthamiana* (Wu et al., 2016; Wu et al., 2017).

**Figure 1.**
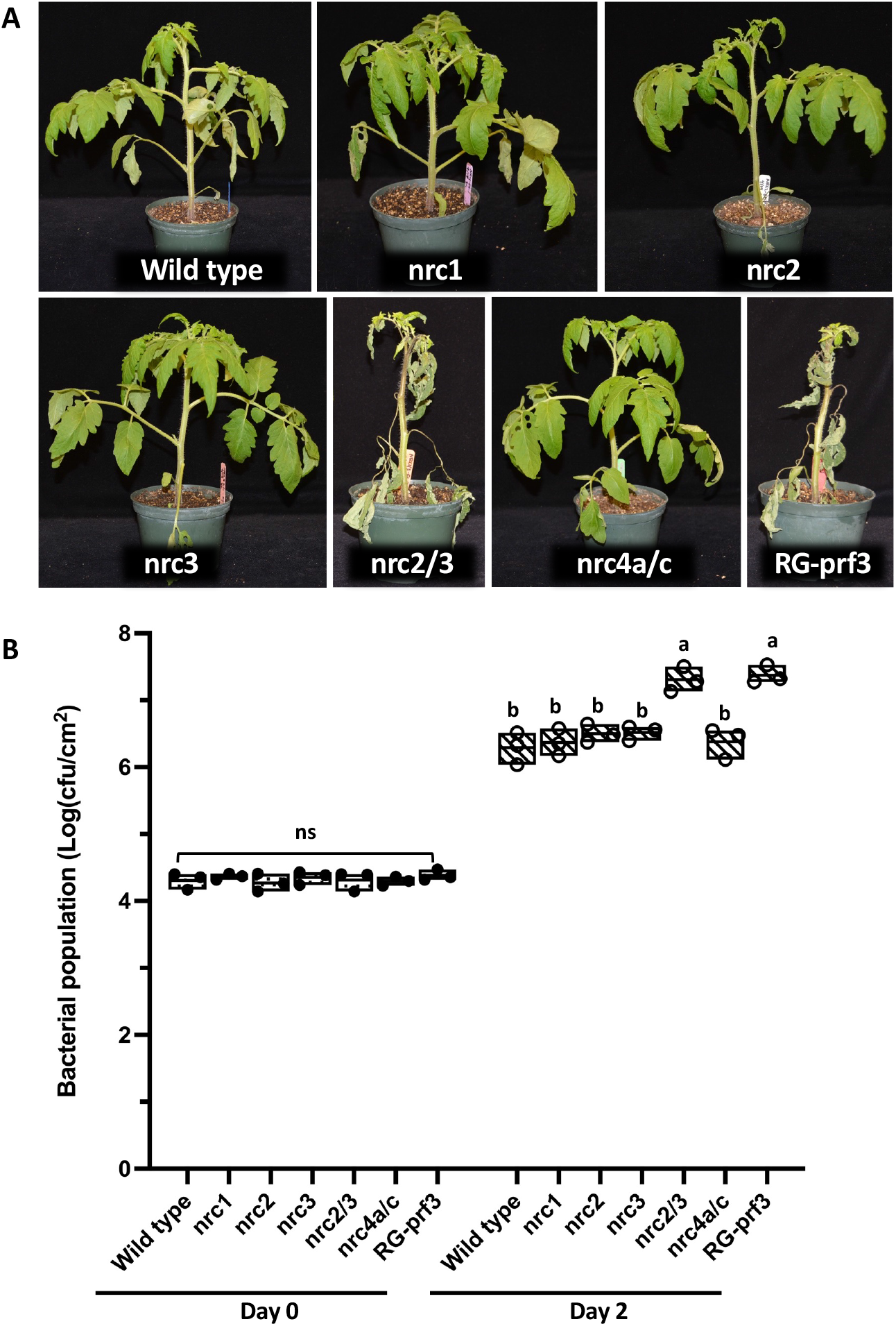
Nrc2 and Nrc3 are required for AvrPto- and AvrPtoB-triggered immunity in tomato. Four-week-old plants of Rio Grande (RG)-PtoR (wild type), nrc1, nrc2, nrc3, nrc2/3, nrc4a/c mutants, and RG-prf3 (a *Prf* mutant) were vacuum-infiltrated with 1 x 10^6^ cfu/mL DC3000. **A**, Photographs of disease symptoms were taken 3 days after infiltration. **B**, Bacterial populations were measured at 3 hours (Day 0) and two days (Day 2) after infiltration. Bars show means ± standard deviation (SD). Different letters indicate significant differences based on a one-way ANOVA followed by Student’s *t* test (p < 0.05). ns, no significant difference. Three plants for each genotype were tested per experiment. The experiment was performed three times with similar results.

As a further test of the contribution of Nrc2 and Nrc3 to the NTI response we used strains of DC3000 that have only AvrPto or AvrPtoB and which consequently might induce a weaker NTI response. The nrc2, nrc3, and nrc2/3 mutants, along with RG-PtoR and RG-prf3 plants, were vacuum infiltrated with either DC3000Δ*avrPto* or DC3000Δ*avrPtoB* at 1×10^6^ cfu/mL (Figs. 2 and 3). Similar to RG-prf3, the nrc2/3 double mutant developed severe disease symptoms when inoculated with either DC3000Δ*avrPto* or DC3000Δ*avrPtoB*. Interestingly, the nrc2 and nrc3 single mutants, developed moderate disease symptoms after inoculation with the two DC3000 strains whereas wild-type RG-PtoR plants were completely resistant to both strains. The nrc3 lines, however, exhibited more pronounced disease symptoms than the nrc2 mutants when inoculated with either strain (Figs. 2 and 3). Consistent with these observations, bacterial populations were about five times higher in the nrc3 mutants than in nrc2 and wild-type RG-PtoR plants and were 10-fold higher in the nrc2/3 mutants and RG-prf3 compared to wild-type plants. Similar results were observed with the other independent nrc2, nrc3 and nrc2/3 mutants when inoculated with DC3000Δ*avrPtoB* (Supplemental Fig. S5). Thus, Nrc2 and Nrc3 are both required for Prf/Pto-mediated NTI and they appear to play an additive role in the immune response induced by these DC3000 strains. Moreover, Nrc3 contributes more strongly than Nrc2 to Prf/Pto-mediated NTI.

**Figure 2.**
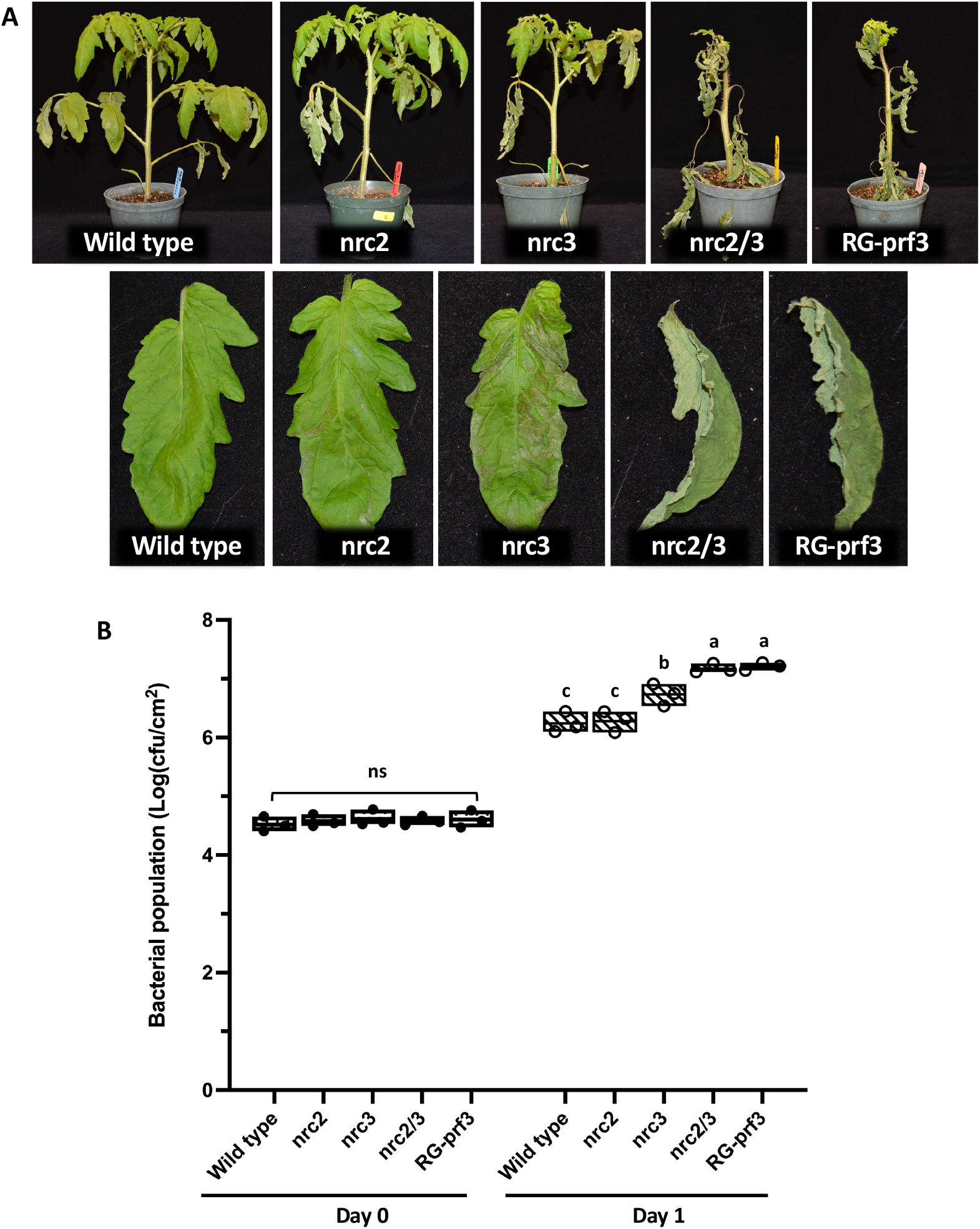
Contribution of Nrc2 and Nrc3 to AvrPtoB-triggered immunity. Four-week-old plants of wild-type RG-PtoR, nrc2, nrc3, nrc2/3, and RG-prf3 plants were vacuum-infiltrated with 1 x 10^6^ cfu/mL DC3000Δ*avrPto*. **A**, Photographs of disease symptoms of the whole plants or leaflets from the 4^th^ leaf were taken 3 days after infiltration. **B**, Bacterial populations were measured 3 hours (Day 0) and one day (Day 1) after infiltration. Bars show means ± SD. Different letters indicate significant differences based on a one-way ANOVA followed by Student’s *t* test (p < 0.05). ns, no significant difference. Three plants for each genotype were tested per experiment. The experiment was performed twice with similar results.

**Figure 3.**
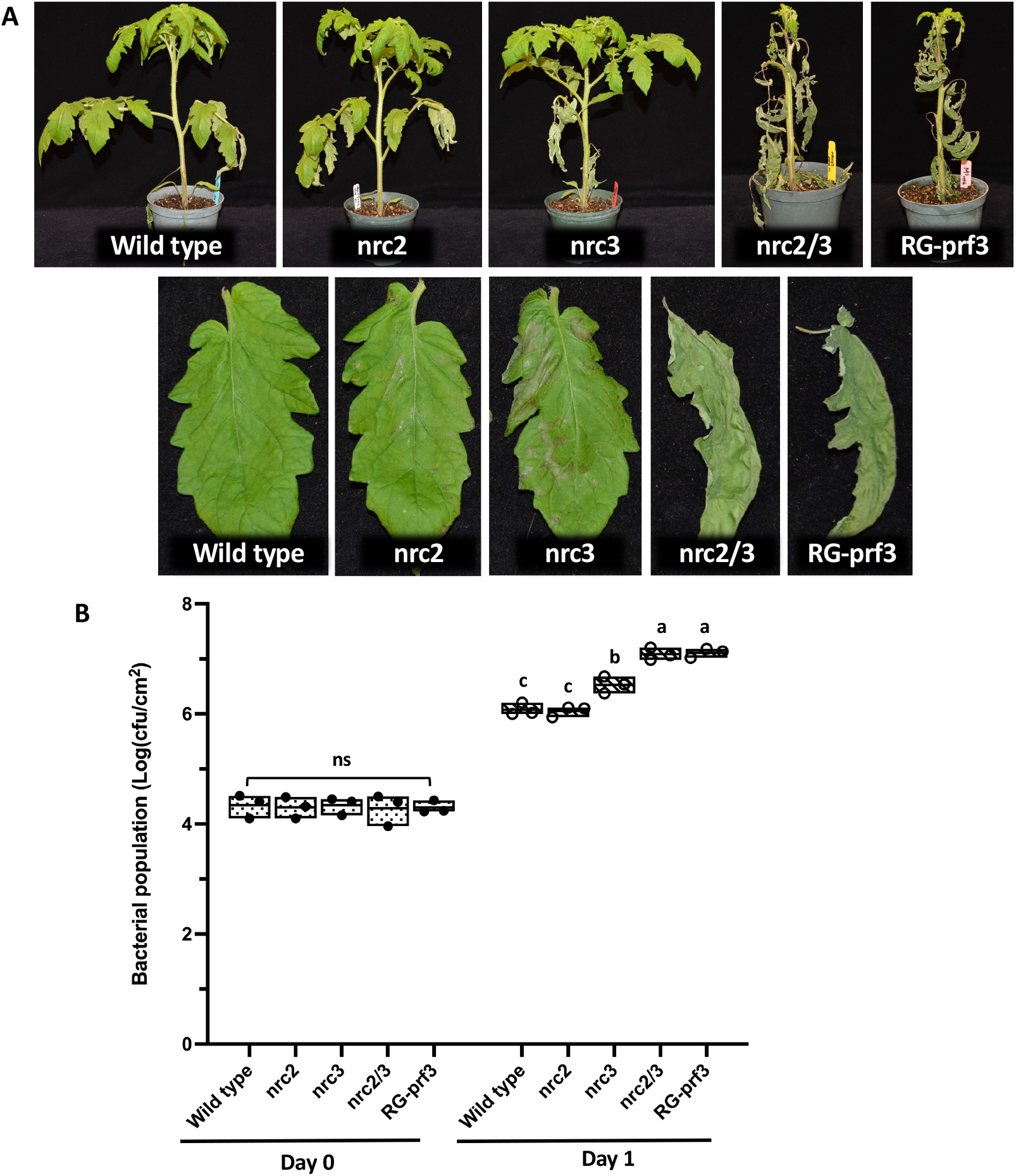
Contribution of Nrc2 and Nrc3 to AvrPto-triggered immunity. Four-week-old plants of wild-type RG-PtoR, nrc2, nrc3, nrc2/3, and RG-prf3 plants were vacuum-infiltrated with 1 x 10^6^ cfu/mL DC3000Δ*avrPtoB*. **A**, Photographs of disease symptoms of the whole plants or leaflets from the 4^th^ leaf were taken at 3 days after inoculation. **B**, Bacterial populations were measured 3 hours (Day 0) and one day (Day 1) after infiltration. Bars show means ± SD. Different letters indicate significant differences based on a one-way ANOVA followed by Student’s *t* test (p < 0.05). ns, no significant difference. Three plants for each genotype were tested per experiment. The experiment was performed twice with similar results.

The additive effect of Nrc2 and Nrc3 was less evident when the plants were inoculated with DC3000Δ*avrPto* or DC3000Δ*avrPtoB* at a lower titer, 1×10^5^ cfu/ml (Supplemental Fig. S6). Specifically, nrc2 plants exhibited disease resistance and supported bacterial growth similar to RG-PtoR, regardless of the inoculating strain. Although nrc3 plants showed more disease lesions compared to RG-PtoR after inoculation with each strain, the bacterial populations in nrc3 were comparable to those in wild-type plants two days post-inoculation.

### Nrc2 and Nrc3 do not contribute to typical PTI responses

Despite employing different activation mechanisms and signaling pathways, PTI and NTI share some common downstream responses and both have been shown in Arabidopsis to be needed for robust immunity (Ngou et al., 2021; Yuan et al., 2021). However, whether *Nrc* genes play a role in PTI responses in tomato is unknown. To explore if any *Nrc* genes contribute to PTI responses against *Pst*, we vacuum-infiltrated the nrc1, nrc2, nrc3, nrc2/3, nrc4a/c and wild-type RG-PtoR plants with the *Pst* strain DC3000Δ*avrPto*Δ*avrPtoB* (DC3000ΔΔ), in which *avrPto* and *avrPtoB* have been deleted and therefore cannot activate NTI (Supplemental Fig. S7A). No significant differences in disease symptoms and bacterial populations were observed between any of the nrc mutants and wild-type plants (Supplemental Fig. S7B). This finding suggests that none of the *Nrc* genes examined, including *Nrc2* and *Nrc3*, play an important role in PTI.

We further investigated if *Nrc2* and *Nrc3* participate in two early PTI-associated responses, generation of reactive oxygen species (ROS) and activation of MAPKs. We exposed the leaves of nrc2, nrc3, nrc2/3 and wild-type RG-PtoR plants to flagellin-derived peptides flg22 or flgII-28 and measured the production of ROS. ROS production was indistinguishable in any of the nrc mutants compared to wild-type plants in response to flg22 or flgII-28 (Supplemental Fig. S7C-D). We also observed no difference between wild-type and any of the nrc mutants for their ability to activate MAPKs in response to flg22 and flgII-28 (Supplemental Fig. S7E). These results indicate that *Nrc2* and *Nrc3* make no demonstrable contribution to these tomato PTI responses against flagellin-associated PAMPs.

### PCD induced by autoactive variants of Prf and Pto requires Nrc2/3 but PCD induced by autoactive M3Kα or Mkk2 does not require Nrc2/3

The observation that the nrc2/3 mutant is completely susceptible to *Pst* DC3000 indicates that *Nrc2* and *Nrc3* play a major role in the Prf/Pto-mediated resistance. To determine the functional position of *Nrc2* and *Nrc3* in the Prf/Pto signaling pathway, we performed a cell death assay in nrc2, nrc3, nrc2/3 and wild-type RG-PtoR plants by expressing in leaves AvrPtoB_1-387_, a variant of AvrPtoB that is a strong Prf/Pto-dependent NTI inducer, or the autoactive variant proteins, Pto^Y207D^, Prf^D1416V^, M3Kα^KD^, or Mkk2^DD^ (Pedley and Martin, 2004; Oh et al., 2010; Du et al., 2012). Upon detection of AvrPto or AvrPtoB, Prf/Pto induce multiple NTI responses including the activation of MAPK cascades, of which M3Kα and Mkk2 are two components (del Pozo et al., 2004; Pedley and Martin, 2004; Oh and Martin, 2011a). Expression in leaves of AvrPtoB_1-387_, Pto^Y207D^, Prf^D1416V^, M3Kα^KD^, or Mkk2^DD^, triggers programmed cell death (PCD) in wild-type RG-PtoR plants (Fig. 4). We observed no decrease in PCD in leaves of the nrc2 or nrc3 plants expressing each of the proteins, as compared to the RG-PtoR leaves (Fig. 4A-C). However, leaves of the nrc2/3 mutant showed compromised PCD upon expression of the proteins AvrPtoB_1-387_, Pto^Y207D^ and Prf^D1416V^ but not of the M3Kα^KD^ and Mkk2^DD^. These observations indicate that Nrc2 and Nrc3 act downstream of or together with the Prf/Pto complex and upstream or independently of MAPK signaling.

**Figure 4.**
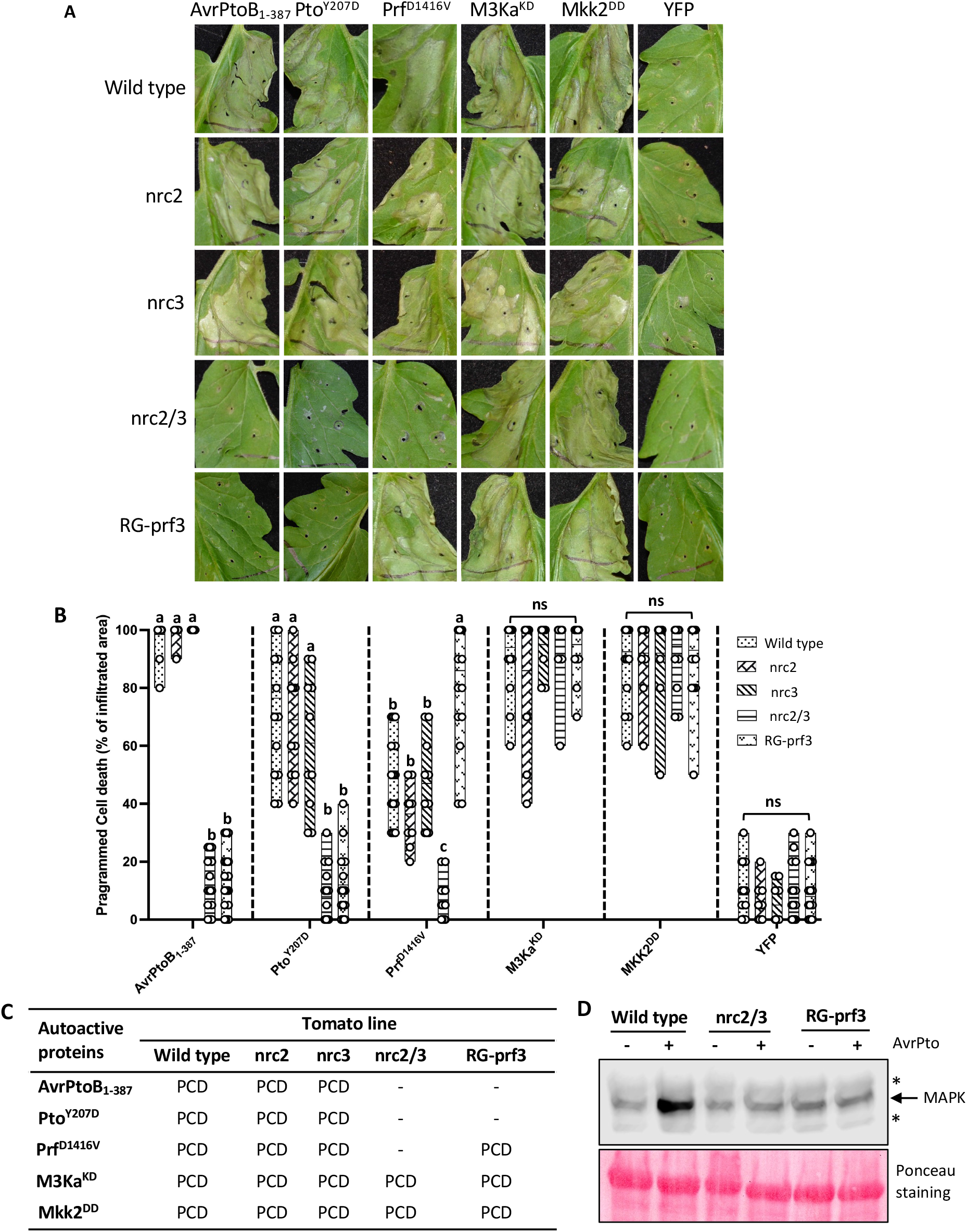
Nrc2 and Nrc3 together contribute Prf/Pto-triggered programmed cell death and MAPK activation in tomato. **A-B**, Leaves of five-week-old plants of wild-type RG-PtoR, nrc2, nrc3, nrc2/3, and RG-prf3 plants were syringe infiltrated with *Agrobacterium tumefaciens* strains carrying constructs of avrPtoB_1-387_, Pto^Y207D^, Prf^D1416V^, M3Ka^KD^, Mkk2^DD^, or YFP (OD_600_ = 0.1 for M3Ka^KD^ and Mkk2^DD^; OD_600_ = 0.3 for the rest). **A**, Photographs 4 days after infiltration. **B**, Average programmed cell death (PCD) scores. Different letters indicate significant differences based on a one-way ANOVA followed by Student’s *t* test (p < 0.05). ns, no significant difference. Results are based on twenty (avrPtoB_1-387_, Pto^Y207D^, Prf^D1416V^ or YFP) or ten (M3Ka^KD^ and Mkk2^DD^) technical replicates from three biological experiments. **C**, Summary of PCD with transient expression of autoactive proteins in tomato leaves of each genotype. Dashes indicate no PCD. **D**, MAPK activation induced by AvrPto. Wild-type RG-PtoR, nrc2/3, and RG-prf3 plants were syringe infiltrated with pER8:EV (empty vector; -) or pER8:avrPto (+). Leaves were treated with 5 µM estradiol and leaf discs were collected 6 hours later. Proteins were extracted from a pool of leaf discs from three plants and subjected to immunoblotting using an anti-pMAPK antibody that detects phosphorylated MAPKs. The photographs shown are derived from the same immunoblot with identical exposure times. Ponceau staining indicates equal loading of protein. Asterisks indicate nonspecific bands. This experiment was performed twice with similar results.

### Nrc2/3 are required for Prf/Pto-mediated activation of MAPKs

To distinguish whether Nrc2/3 act upstream or independently of MAPK signaling we used an estradiol-inducible AvrPto construct (pER8:AvrPto) developed previously (Pedley and Martin, 2004). Leaves of RG-PtoR, RG-prf3, and nrc2/3 were agroinfiltrated with the pER8:AvrPto construct and treated with estradiol one day after agroinfiltration. Six hours later proteins were extracted and activation of MAPKs was tested by immunoblotting using anti-pMAPK antibody (Zhang et al., 2020). With these conditions, we observed an increase in MAPK activity in the wild-type Prf/Pto-expressing RG-PtoR plants. No increase in MAPK activity was detected in RG-prf3 in response to expression of AvrPto due to the non-functional Prf/Pto pathway in these plants. Notably, MAPK activation triggered by expression of AvrPto was completely abolished in the *nrc2*/3 mutant (Fig. 4D). These results indicate that Nrc2/3 are required for downstream activation of MAPK signaling that is induced by the Prf/Pto pathway and that MAPK cascades play a key role in activating Prf/Pto-mediated NTI responses.

### PCD induced by an autoactive Nrc3 variant requires Prf

Next, to better understand the relationship of Nrc2/3 and Prf/Pto in the signaling pathway, we generated a constitutively-active variant of Nrc3 by creating amino acid substitutions in the methionine-histidine-aspartate (MHD) motif resulting in Nrc3^H478AD479V^ (Fig. 5A; (Sueldo et al., 2015)). These substitutions in the MHD motif of the Nrc3 protein had no observable effect in protein stability (Fig. 5B). While transient expression of the Nrc3^H478AD479V^ protein failed to cause cell death in tomato leaves, it did trigger a strong PCD in leaves of wild-type *N*. *benthamiana* (Nb1) plants (Fig. 5C). This allowed us to examine the possible requirement of Prf for Nrc3 function by expressing the autoactive Nrc3 in leaves of an *N. benthamiana* line that carries a hairpin-Prf insertion (hpPrf) that decreases expression of Prf (Fig. 5C). Interestingly, expression of the constitutively-active form of the Nrc3 protein did not trigger PCD in the hpPrf line. These results indicate that Nrc3, and probably Nrc2, are dependent on Prf to trigger the downstream programmed cell death during NTI responses.

**Figure 5.**
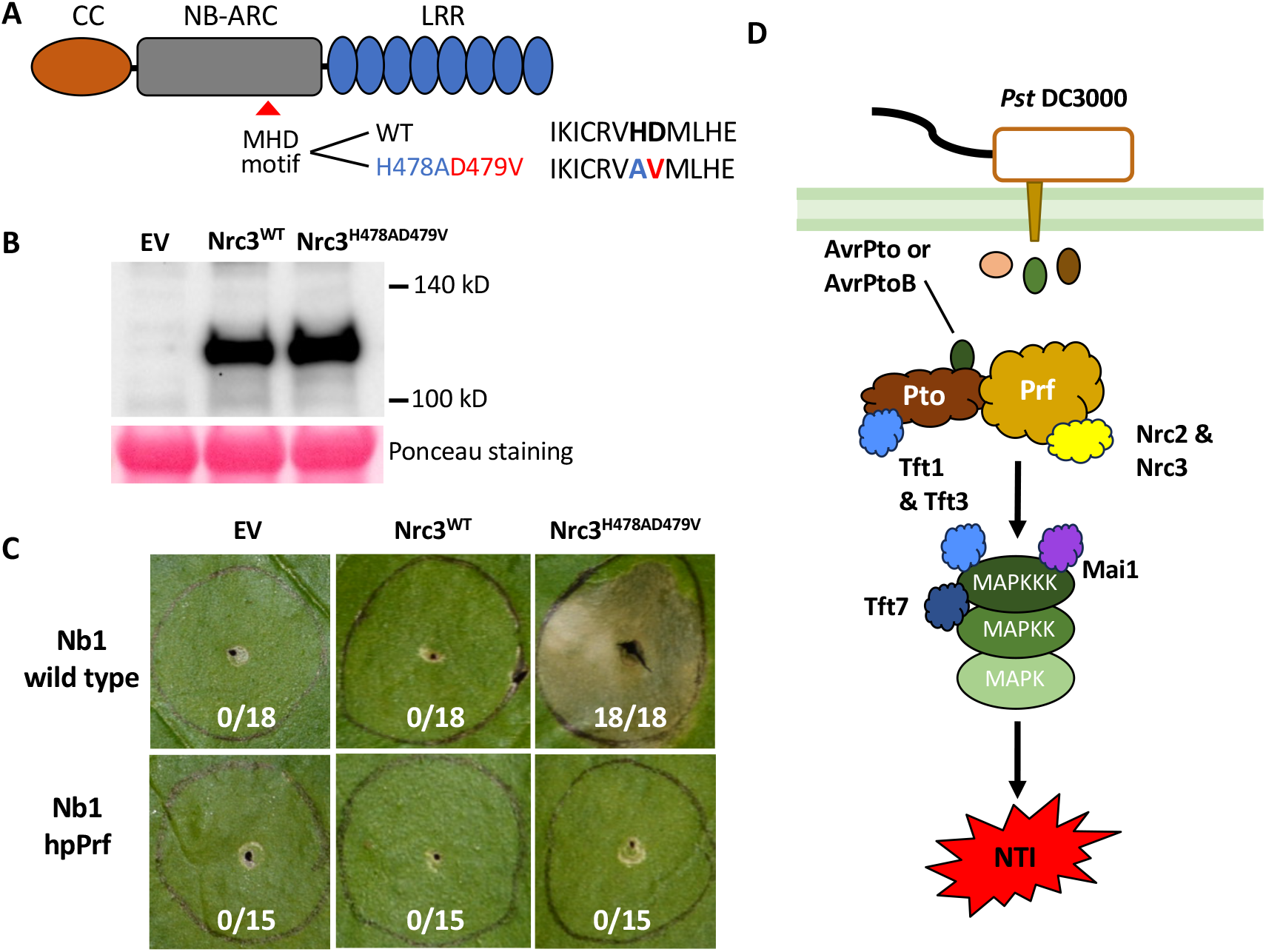
A constitutive-active variant of Nrc3 requires Prf to induce cell death in *Nicotiana benthamiana*. **A**, Schematic representation of mutagenesis of the MHD motif in the Nrc3 protein. **B**-**C**, The GV2260 strain expressing either the wild-type Nrc3, the constitutive-active variant Nrc3^H478AD479V^, or the control (empty vector; EV) with OD=0.3 into the leaves of 5-week-old *Nicotiana benthamiana* Nb1 wildtype and hpPrf plants. Proteins were extracted from *N. benthamiana* Nb1 wild-type leaves expressing either EV, Nrc3^WT^, or Nrc3^H478AD479V^ and detected by immunoblotting with the ⍺-HA antibody (**B**). Photographs were taken 6 days after infiltration (**C**). Shown are the number of times cell death was observed over the total number of agroinfiltrations performed. This experiment was repeated twice with similar results. **D**, Schematic drawing showing Nrc2 and Nrc3 act co-dependently with Prf and induce MAPK signaling and possibly other Prf/Pto-mediated immune responses in tomato (see Discussion for additional details).

### Nrc2 and Nrc3 do not play a role in Ptr1-associated PCD

The CC-NLR-encoding *Ptr1* gene, which was identified recently in a wild relative of tomato, *Solanum lycopersicoides*, confers resistance to *Pst*, *Xanthomonas* spp. and *Ralstonia pseudosolanacearum* by detecting multiple sequence-diverse type III effectors, including AvrRpt2 (Mazo-Molina et al., 2019; Mazo-Molina et al., 2020; Kim et al., 2022; Ahn et al., 2023b; Kim et al., 2023; Tsakiri et al., 2023). It is unknown whether any Nrc protein is required for Ptr1-mediated immunity. To address this question, we carried out a cell death assay by co-expressing AvrRpt2 and Ptr1 in the leaves of nrc1, nrc2/3, nrc4a/c and wild-type RG-PtoR plants. Strong cell death was observed in the wild-type plants, as well as in nrc1, nrc2/3 and nrc4a/c mutants four days after infiltration (Supplemental Fig. S8). This suggests that Nrc1, Nrc2, Nrc3, and Nrc4a/c are not essential for the Ptr1-mediated immune response in tomato.

### Nrc2 and Nrc3 play a role in resistance to some *Pst* strains that lack AvrPto and AvrPtoB

Prf is the only NLR protein in tomato that is known to act with Nrc2 and Nrc3 to mediate resistance to *Pst* DC3000. To investigate whether Nrc2 and Nrc3 contribute to other R gene-mediated immune responses, we vacuum-infiltrated nrc2/3 and wild-type RG-PtoR plants with the *Pst* strains DC3000Δ*avrPto*Δ*avrPtoB*Δ*fliC* (DC3000ΔΔΔ), T1Δ*avrPtoB*Δ*fliC* (T1ΔΔ) (Almeida et al., 2009), or Pto19Δ*avrPtoB* (Pto19Δ) (Kunkeaw et al., 2010), in which *avrPto* and/or *avrPtoB* are absent or deleted and therefore cannot activate Prf/Pto-mediated NTI. Two days after inoculation, the nrc2/3 mutant displayed similar disease symptoms as wild-type RG-PtoR, when inoculated with DC3000ΔΔΔΔ but enhanced susceptibility when inoculated with either T1ΔΔ or Pto19Δ (Fig. 6). The bacterial population in the nrc2/3 plants was similar to that in RG-PtoR when inoculated with DC3000ΔΔΔ or T1ΔΔ but was significantly higher than that in RG-PtoR plants when inoculated with Pto19Δ. Together, these observations suggest that Nrc2 and Nrc3 might act with another NLR sensor of an effector present in Pto19 and T1. We therefore generated a genome sequence for Pto19 and compared its type III effector repertoire with the known repertoires of T1 and DC3000 (Supplemental Fig. S9). This analysis revealed that Pto19 and T1 have eight effectors in common that are not present in DC3000. We tested whether Nrc2 and Nrc3 mediate recognition of these effectors by inoculating previously developed DC3000Δ*avrPto*Δ*avrPtoB* strains expressing each effector onto nrc2/3 and wild-type plants (Mazo-Molina et al., 2019). None of the plants showed differential symptoms or altered bacterial growth (note that a DC3000 strain carrying HopAI1 was not available, and the one carrying HopAS1 appeared to be recognized by both wild type and nrc2/3 plants and its analysis will be presented separately). It remains possible that Nrc2 and Nrc3 mediate recognition of HopAI1 or an allelic variant of an effector present in T1 and Pto19 that is different in DC3000, or that an effector in DC3000 suppresses recognition of an effector present in all these strains and these possibilities will be investigated in the future.

**Figure 6.**
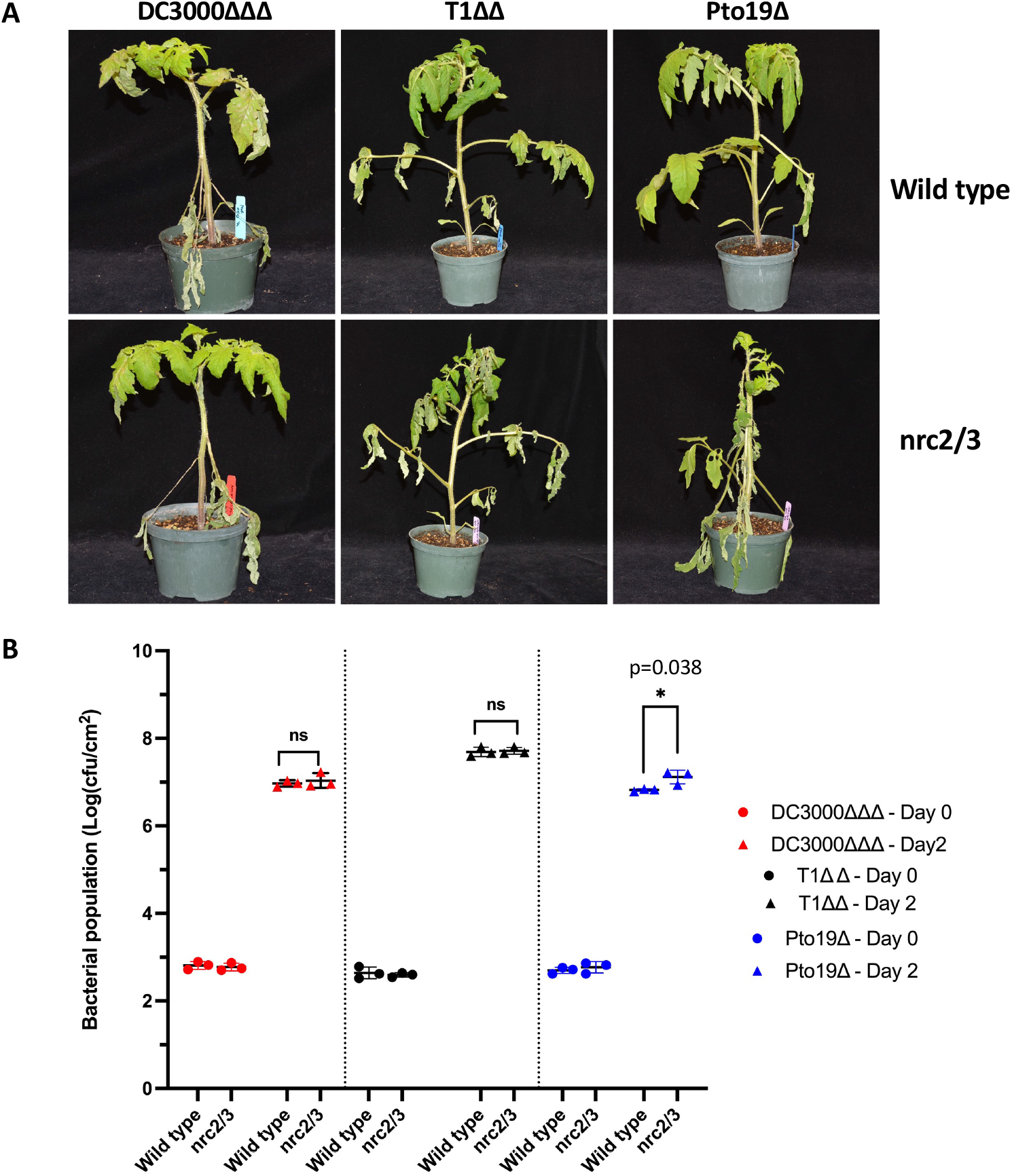
Nrc2 and Nrc3 probably contribute to other R gene-mediated resistance in tomato to *Pseudomonas syringae* pv. *tomato*. **A**, Four-week-old plants of wild-type RG-PtoR and nrc2/3 plants were vacuum-infiltrated with 2 x 10^4^ cfu/mL DC3000Δ*avrPto*Δ*avrPtoB*Δ*fliC* (DC3000ΔΔΔ), 3 x 10^4^ cfu/mL T1Δ*avrPtoB*Δ*fliC* (T1ΔΔ) or 2 x 10^4^ cfu/mL Pto19Δ*avrPtoB* (Pto19Δ). Photographs of disease symptoms of the whole plants were taken 4 days after inoculation. **B**, Bacterial populations were measured 3 hours (Day 0) and one day (Day 2) after infiltration. Bars show means ± SD and the middle horizontal line is the mean of the three plants. Statistical significance was determined by pairwise t test (* p <0.05; ns, no significance). Three plants for each genotype were tested per experiment. The experiment was performed twice with similar results.

## Discussion

The tomato *Nrc2* and *Nrc3* genes were originally identified as being functionally redundant, playing key roles in Prf/Pto-mediated cell death and resistance against *P. syringae* pv. *tomato* (*Pst*) (Wu et al., 2016; Wu and Kamoun, 2021). When the expression of *Nrc2* and *Nrc3* was reduced by virus-induced gene silencing (VIGS) in tomato, the Prf/Pto-mediated cell death was abolished but disease resistance to *Pst* expressing *avrPto* and *avrPtoB* was only partially affected (Wu and Kamoun, 2021). The implication of these results might be limited by the insufficient silencing of the *Nrc2* and *Nrc3* genes by VIGS, a shortcoming of this technique (Rosli et al., 2013). Therefore, the extent to which a complete knockout of the *Nrc2* and *Nrc3* genes might impact Prf/Pto-mediated disease resistance to *Pst* remained unclear. Here we have developed CRISPR/Cas9-mediated loss-of-function nrc mutants in tomato and used them to investigate the contributions of *Nrc* genes, particularly *Nrc2* and *Nrc3*, to the Prf/Pto-mediated hypersensitive response (HR)-associated cell death and disease resistance to *Pst* in tomato. Our results show that Nrc2 and Nrc3 appear to function redundantly in their response to *Pst* DC3000 but can be seen to function additively in response to DC3000 strains that elicit a weaker NTI response. In addition, our results indicate that Nrc3 acts co-dependently with Prf and upstream of MAPK signaling, that Nrc2 and Nrc3 are not required for the HR-associated cell death triggered by the Ptr1 sensor NLR, and that they may act with another unknown sensor NLR in tomato to mediate resistance to certain strains of *Pst* lacking AvrPto and AvrPtoB.

Previous studies have shown that Nrc2 and Nrc3, paired with various sensor NLRs, exhibit redundant functionality that confers immunity to certain bacteria, viruses, and oomycetes (Wu et al., 2016; Wu et al., 2017; Wu and Kamoun, 2021). Here, loss-of-function mutation of either *Nrc2* or *Nrc3* alone did not compromise resistance in tomato to *Pst* DC3000 which has AvrPto and AvrPtoB. In addition, the single mutants nrc2 and nrc3 still developed strong cell death triggered by AvrPtoB_1-387_ or autoactive variants of Pto or Prf, but the double mutant nrc2/3 did not develop cell death induced by these proteins. These observations support the notion that Nrc2 and Nrc3 can act redundantly in Prf/Pto-mediated NTI in tomato. However, unlike the previous finding that silencing of both *Nrc2* and *Nrc3* partially compromised disease resistance to *Pst* in tomato (Wu and Kamoun, 2021), we observed that loss-of-function mutations in both *Nrc2* and *Nrc3* led to complete susceptibility, comparable to the RG-prf3 which contains a mutation in *Prf* that makes the Prf/Pto pathway nonfunctional. This indicates that Nrc2 and Nrc3 play overlapping important roles in the Prf/Pto pathway in tomato.

Interestingly, while exhibiting redundancy in response to DC3000, Nrc2 and Nrc3 also appear to act additively in the Prf/Pto-mediated NTI pathway. This was observed in inoculations of DC3000 variants lacking either *avrPto* or *avrPtoB* on the single mutants nrc2 and nrc3, particularly nrc3, in which case more pronounced disease symptoms developed compared to the completely resistance control RG-PtoR. These results indicate that Nrc3 plays a stronger role than Nrc2 in disease resistance to *Pst*. Consistent with our findings, tomato Nrc3 could partially restore Pto-mediated cell death but Nrc2 did so only weakly in *Nrc2/3-*silenced *N. benthamiana* (Wu et al., 2016; Wu et al., 2017). Similarly, in *N. benthamiana* Nrc3 is primarily responsible for mediating the hypersensitive cell death caused by the cell-surface receptor Cf4 (Nrc2 only partially complemented Cf4/Avr4-triggered cell death as compared to Nrc3) (Kourelis et al., 2022). The predominant role of tomato Nrc3 in Prf/Pto-mediated NTI responses might be due to a greater accumulation of Nrc3 protein corresponding to the higher transcript abundance of *Nrc3* compared to *Nrc2*, under both unchallenged and *Pst*-challenged conditions (Supplemental Fig. S1 and Supplemental Table S1). Consequently, when *Nrc3* is knocked out, the remaining Nrc2 protein might be insufficient to promote full Prf/Pto downstream signaling. Alternatively, Nrc3 might have greater stability or higher binding affinity for the Prf/Pto complex or another interacting protein, thereby playing a more significant role than Nrc2 during NTI responses. That hypothesis is supported by the observations that the additive effect of Nrc2 and Nrc3 in tomato immunity against *Pst* was bacterial dose-dependent, that is, Nrc2 or Nrc3 alone could activate a robust Prf/Pto-mediated NTI response when the bacterial population was relatively low (Supplemental Fig. S6).

The nrc2/3 mutant was compromised in PCD in response to AvrPtoB_1-387_ or autoactive variants of Pto and Prf, and the *N*. *benthamiana* hpPrf line also was compromised in PCD in response to a constitutively-active variant of Nrc3. Together, these results demonstrate the essential role of Nrc2 and Nrc3 proteins in Prf/Pto-triggered PCD, and are also consistent with the finding that Nrc2 and Nrc3 directly interact with the Prf/Pto complex prior to its activation (Sheikh et al., 2023). Importantly, our results extend the previous observations that Prf requires Nrc2 and Nrc3 for immunity to the finding that Nrc3 (and likely Nrc2) also requires Prf for triggering the immune response. Our study also bridges a gap by elucidating the connection between the helper NLRs Nrc2/3 and MAPK signaling. While the significance of activation of mitogen-activated protein kinases (MAPKs) for NTI responses in Arabidopsis and tomato has been well-established (del Pozo et al., 2004; Oh and Martin, 2011b; Su et al., 2017; Su et al., 2018; Roberts et al., 2019; Zhang and Zhang, 2022), the precise roles of Nrc2 and Nrc3 in MAPK signaling during NTI responses in tomato remains elusive. Our study reveals that Nrc2 and Nrc3 proteins are required for triggering downstream NTI-associated MAPK signaling. Based on these observations, we propose a model for the role of Nrc2 and Nrc3 in the Prf/Pto-mediated NTI response (Fig. 5D). In this model, Nrc2 and Nrc3 act co-dependently with the Prf/Pto complex and induce the downstream MAPK signaling leading to PCD, a key feature of NTI responses resulting in disease resistance.

Recently, Sheikh et al. (2023) also proposed a model for how Nrcs are involved in the Prf/Pto-mediated immunity. In their model, the Prf/Pto complex, comprised of Prf, Pto, Nrc2/3, Tft1, and Tft3, dissociates into two distinct NTI signaling modules following effector recognition. The Tft3-dependent branch is required for full activation of MAPK cascades, and partially contributes to PCD, while the Nrc2/3-dependent branch is essential for full PCD formation and restriction of pathogen growth. Specifically, in their model, Nrc proteins, including Nrc2 and Nrc3, are not required for downstream MAPK signaling during NTI responses, based on the observations that Nrc2 and Nrc3 had no interaction with Tft3 or M3Kα in co-immunoprecipitation experiments (Sheikh et al., 2023). While our model mostly aligns with theirs, a key difference lies in the contribution of Nrc2/3 to the Tft3-mediated programmed cell death and MAPK activation. Our findings that Prf and Nrc3 (and likely Nrc2) act co-dependently to induce programmed cell death and Nrc2 and Nrc3 are required to activate downstream MAPK signaling support the notion that Nrc2/3 and Prf/Pto form an integral complex and together participate in any downstream NTI signaling pathway that leads to PCD, likely including the Tft3-dependent signaling pathway that partially contributes to PCD, and the Tft3-dependent activation of MAPK cascades, as reported in Sheikh et al. (2023).

Further research is needed to understand how Prf/Pto activation is transmitted to Nrc2 and Nrc3, identify the downstream targets modulated by these helper NLRs, and establish the missing molecular link between Nrc2/3 and its interacting proteins. Several genes have been identified to play key roles in Prf/Pto-mediated cell death and disease resistance against *Pst* in tomato. The 14-3-3 proteins, Tft1 and Tft3, have been recently identified as interacting partners of Pto and the protein kinase M3Kα (Sheikh et al., 2023). Another 14-3-3 protein, Tft7, interacts with M3Kα and Mkk2 and positively regulates Prf/Pto-mediated PCD in tomato (Oh et al., 2010; Oh and Martin, 2011b). The tomato M3Kα-interacting protein, Mai1, a receptor-like cytoplasmic kinase, acts upstream of M3Kα and positively contributes to PCD and disease resistance associated with Prf/Pto activation (Roberts et al., 2019). However, it remains unknown whether Nrc2 and Nrc3 interact with or regulate the immune responses mediated by Mai1, Tft1, Tft3, or Tft7. Future investigation of these regulatory networks will provide valuable insights into the mechanisms by which helper Nrcs function in conjunction with sensor NLRs and shed light on the intricate interactions and signaling pathways involved in the NTI response in plants.

## Materials and Methods

### Generation of tomato nrc2/3 mutants using CRISPR/Cas9

To generate nrc1, nrc2, nrc3, and nrc4a/c mutants in the tomato cultivar Rio Grande (RG)-PtoR, which has the *Pto* and *Prf* genes, we designed guide RNAs (gRNAs; Supplemental Fig. S2) targeting the first or second exons of each *Nrc* gene using the software Geneious R11 (Kearse et al., 2012). Each gRNA cassette was cloned into the p201N:Cas9 binary vector as described previously (Jacobs et al., 2017). To mutate both *Nrc4a* and *Nrc4c* in the same plant, gRNAs targeting *Nrc4a* and *Nrc4c* were cloned into the binary vector for transformation. Tomato transformation was performed at Center for Plant Biotechnology Research at the Boyce Thompson Institute as described previously (Zhang et al., 2020). Mutations were confirmed by Sanger sequencing at the Biotechnology Resource Center (BRC) at Cornell University (Supplemental Table S2). The two nrc2/3 mutant lines were generated by crossing nrc2 or nrc2-2 to nrc3. F1s were then selfed and F2 populations were genotyped to screen for homozygous mutations in both the *Nrc2* and *Nrc3* genes.

### Bacterial strains

*Pseudomonas syringae* pv. *tomato* (*Pst*) stains DC3000 (Buell et al., 2003), DC3000Δ*avrPto* (Ronald et al., 1992), DC3000Δ*avrPtoB* (Lin and Martin, 2005), DC3000Δ*avrPto*Δ*avrPtoB* (Lin and Martin, 2005), DC3000Δ*avrPto*Δ*avrPtoB*Δ*fliC* (gift from Alan Collmer, Cornell U), T1Δ*avrPtoB*Δ*fliC* (Almeida et al., 2009), and Pto19Δ*avrPtoB* (Kunkeaw et al., 2010) were grown on King’s B (KB) semi-selective media with appropriate antibiotics at 30°C (Supplemental Table S3). *Agrobacterium tumefaciens* strains GV2260 and 1D1249 were grown on LB media with appropriate antibiotics at 30°C (Supplemental Table S4).

### Bacterial inoculation

Four-week-old plants of nrc mutants and wild-type RG-PtoR were vacuum-infiltrated with various *Pst* strains, including DC3000, DC3000Δ*avrPto*, or DC3000Δ*avrPtoB* at 1 x 10^6^ cfu/mL, DC3000Δ*avrPto*Δ*avrPtoB* at 5 x 10^4^ cfu/mL, DC3000Δ*avrPto*Δ*avrPtoB*Δ*fliC* at 2 x 10^4^ cfu/mL, T1Δ*avrPtoB*Δ*fliC* at 3 x 10^4^ cfu/mL, and Pto19Δ*avrPtoB* at 2 x 10^4^ cfu/mL. Bacterial populations were measured at 3 h (Day 0) and one or two days after inoculation (Day 1 or Day 2). Photographs of disease symptoms were taken three to six days after bacterial inoculation.

### ROS assay

Production of reactive oxygen species (ROS) was measured as described previously (Clarke et al., 2013; Zhang et al., 2022). In brief, leaf discs from 5-week-old tomato plants were collected and floated in water overnight. Water was then removed and replaced with a solution containing 34 µg/mL luminol (Sigma-Aldrich) and 20 µg/mL horseradish peroxidase, in combination with 50 nM flg22 or 50 nM flgII-28. ROS production was measured using a Synergy 2 microplate reader (BioTek).

### MAPK phosphorylation assay

To test flagellin-induced MAPK activation, six leaf discs of nrc2, nrc3, or nrc2/3 mutants and wild-type RG-PtoR plants were floated in water overnight. The leaf discs were then incubated with 20 nM flg22, 25 nM flgII-28, or water for 10 min, and frozen immediately in liquid nitrogen. To test AvrPto-induced MAPK activation, wild-type RG-PtoR and nrc2/3 mutants were syringe infiltrated with *Agrobacterium tumefaciens* GV2260 expressing pER8:AvrPto or pER8:EV (empty vector) (OD=0.3) as the control. One day later, leaves were sprayed with 5 μM estradiol and leaf discs were collected six hours later and frozen immediately in liquid nitrogen. Protein was extracted using a buffer containing 50 mM Tris-HCl (pH 7.5), 10% glycerol, 2 mM EDTA, 1% Triton X-100, 5 mM DTT, 1% protease inhibitor cocktail (Sigma-Aldrich), 0.5% phosphatase inhibitor cocktail 2 (Sigma-Aldrich). MAPK phosphorylation was determined using an anti-phospho-p44/42 MAPK (Erk1/2) antibody (anti-pMAPK; Cell Signaling).

### *Agrobacterium*-mediated transient protein expression and cell death assays

*Agrobacterium tumefaciens* strains GV2260 and 1D1249 harboring the noted expression vectors were grown in YFP media with appropriate antibiotics at 30°C overnight. *Agrobacterium* cells were induced using the AGROBEST protocol (Wu et al., 2014) or with infiltration buffer (10 mM MgCl_2_, 10 mM morpholineethane sulfonic acid [MES, pH 5.6], and 200 mM acetosyringone) for 4 h at room temperature. The bacterial culture was then centrifuged, washed, resuspended in infiltration buffer and adjusted to the final indicated OD_600_. The third leaves of five-week-old tomato plants or *Nicotiana benthamiana* were syringe infiltrated and the whole plants were placed in a growth chamber (24°C/day and 22°C/night). Photographs of cell death in the infiltrated areas were taken 4 to 6 days after infiltration. Cell death was scored as the percentage of dead area within the infiltrated leaf area.

### Generation of the Nrc3 autoactive variant

The coding region of the tomato *Nrc3* gene was amplified from tomato cDNA using Phusion Hot Start II DNA polymerase (ThermoFisher Scientific) and gene-specific primers (Supplemental Table S2). Autoactive mutant of Nrc3 (Nrc3^H478AD479V^) was generated by introducing codon changes to make a histidine (H) to alanine (A) substitution and an aspartic acid (D) to valine (V) substitution in the MHD motif of Nrc3 based on the NbNrc3 autoactive mutant reported previously (Derevnina et al., 2021). To generate this variant, a 249 bp gBlcok containing the mutated DNA sequence was synthesized (Integrated DNA technology; Supplemental Table S2) and cloned into pJLSmart along with other coding region of the *Nrc3* gene by Gibson assembly. The gene expression cassette in pJLSmart was then cloned into the destiny vector pGWB414 via recombination reactions using LR Clonase II (ThermoFisher Scientific). Constructs were confirmed by Sanger sequencing and then transformed into *Agrobacterium* strain GV2260 for transient expression in *N. benthamiana*.

### Protein immunoblotting

Total protein was extracted from *N. benthamiana* leaves using 250 μl extraction buffer consisting of 62.5 mM Tris-HCl (pH 6.8), 2% SDS (v/v), 10% glycerol and 5% β-mercaptoethanol. A 12 μL soluble protein solution mixed with 4x Laemmli sample buffer were boiled at 95°C for 5 min before loaded for gel electrophoresis. Protein was loaded on 4% −20% precast SDS-PAGE gel (Bio-Rad), blotted on PVDF membrane (Merck Millipore), inoculated with α-HA primary antibody (1:2000; v/v) and α-rabbit-HRP secondary antibody (1:10000; v/v), and developed with Immobilon forte Western HRP substrate (Millipore Sigma) for 5 min.

### Genome sequencing and effector prediction of *Pst* Pto19

Genomic DNA of *Pst* Pto19 was extracted using the Quick-DNA Miniprep Plus Kit, according to the manufacturer’s instructions (Zymo Research). The concentration of the genomic DNA was adjusted to 100 ng/μL using Qubit dsDNA HS assay (ThermoFisher Scientific). Genomic sequencing was performed using Oxford Nanopore and the complete genome was annotated by Plasmidsaurus (https://www.plasmidsaurus.com/). A database previously generated which contains 14,613 type III effector protein sequences (Dillon et al., 2019) was used in a blastp analysis using the protein sequences of Pto19 as queries, as described previously (Gonzalez-Tobon et al., 2023). Only hits with E values over a significant threshold of <1e-24 were considered significant.

## Acknowledgments

We thank Liam Cleary, Brian Bell, Jay Miller, and Nick Vail for plant care, Joyce Van Eck and Tish Keen for tomato transformation, and Paul Stodghill for Pto19 genome analysis. Funding was provided by National Science Foundation grant IOS-1546625 (GBM).

## Author contributions

NZ and GBM conceived and designed the experiments. NZ designed gRNAs and constructed vectors. NZ and JG performed genotyping experiments, JGT and MF identified and annotated the type III effectors for *Pst* Pto19, NZ, LC, and JG performed phenotyping experiments and analyzed the data. NZ and GBM interpreted the data and wrote the manuscript. All the authors read and approved the manuscript.

## Competing interests

The authors declare that they have no conflict of interest.

## Supplemental Information

**Supplemental Figure S1.**
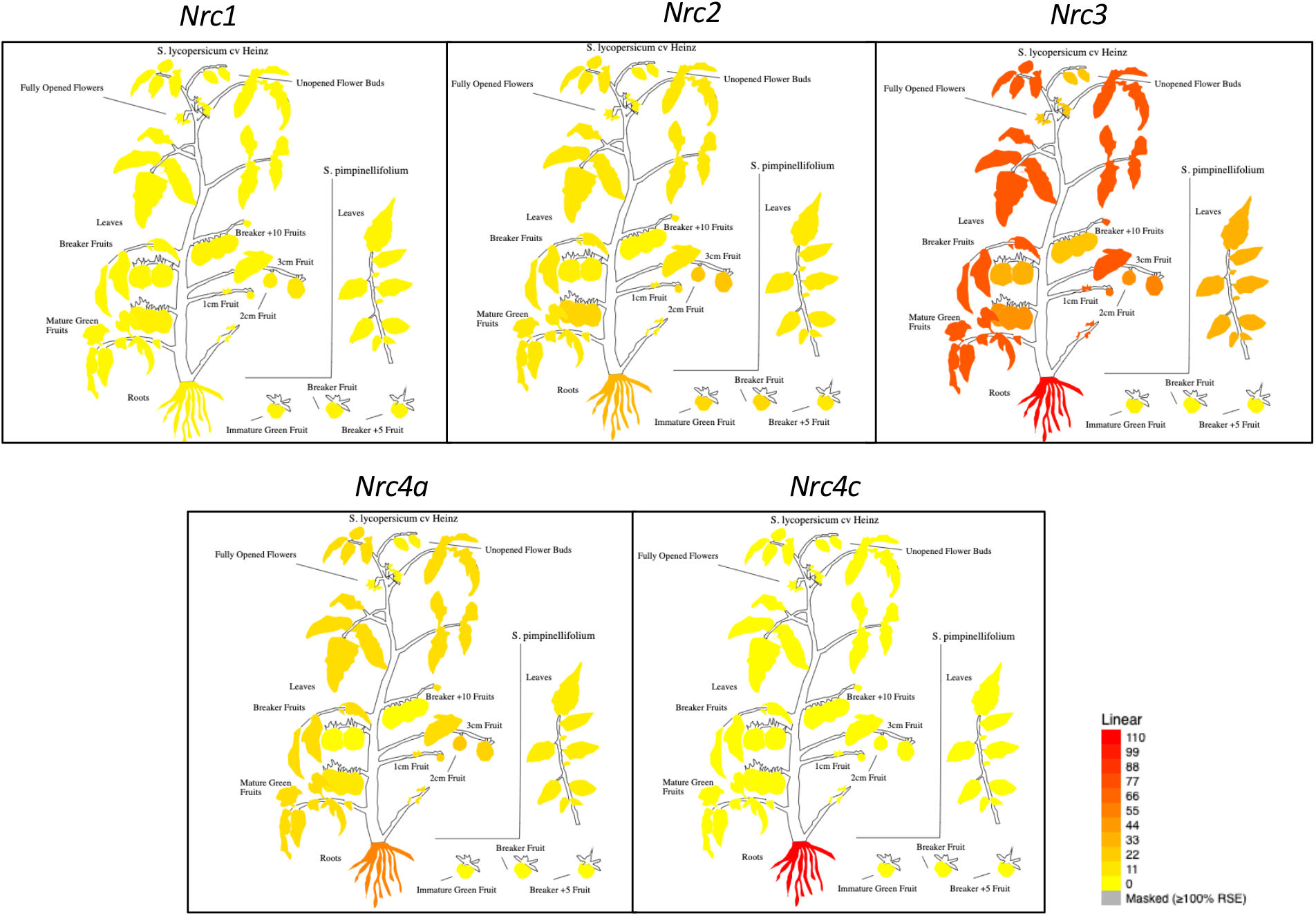
Expression patterns of *Nrc* genes in different tissues/organs of wild-type tomato plants. The figures were generated using the ePlant Tomato Tool (bar.utoronto.ca/eplant_tomato/).

**Supplemental Figure S2.**
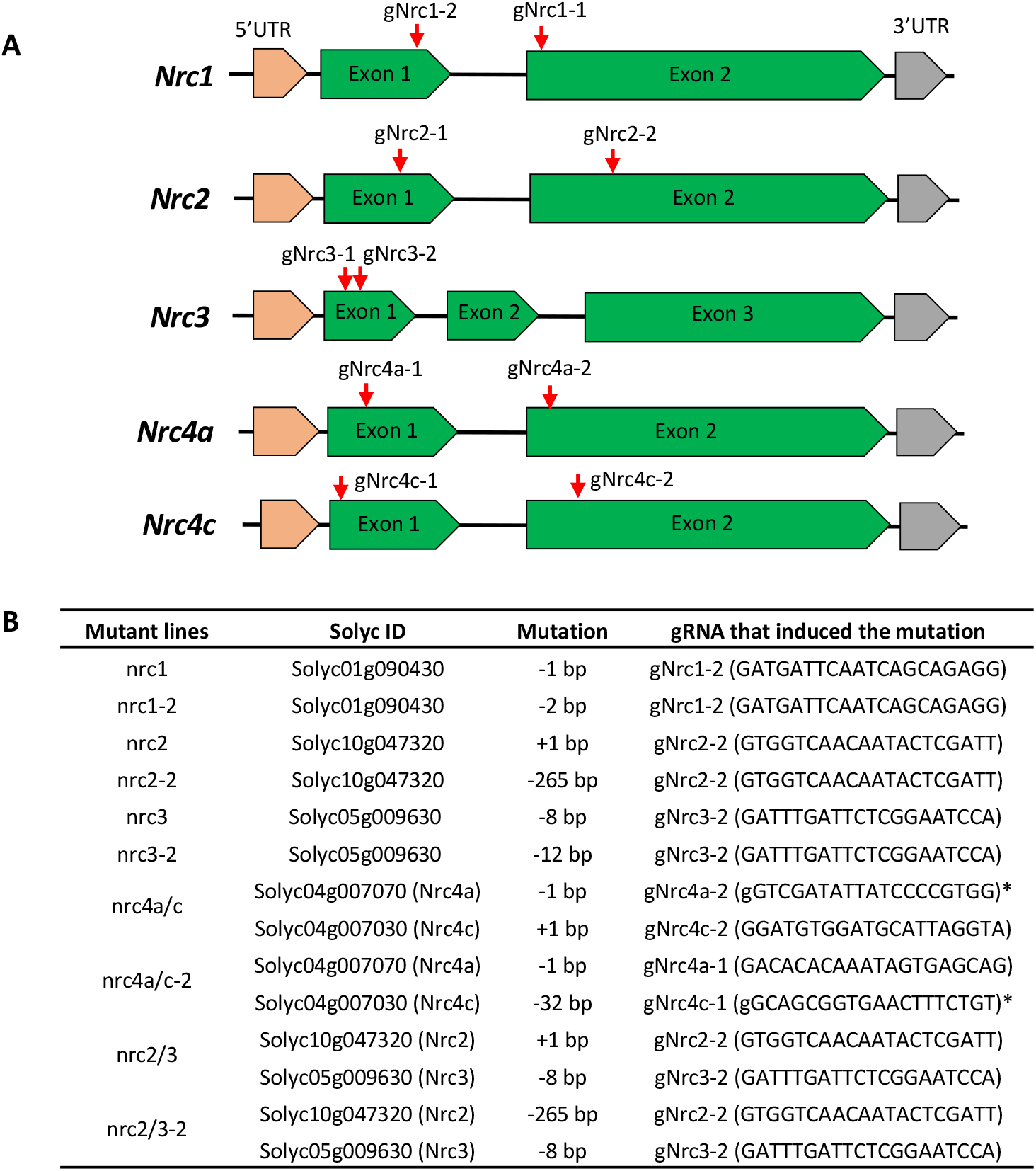
Generation of tomato nrc mutants by CRISPR/Cas9. **A**, Schematics showing the guide-RNA (gRNA) target sites in each *Nrc* gene (gNrcx-x). **B**, Summary of mutations in the nrc mutant plants used in this study. * “g” in lower case means the first nucleotide of the gRNA sequence is not a “G” but was converted to that nucleotide to accommodate the transcription initiation requirement of the U6 promoter. ^#^ The two nrc2/3 mutant lines were generated by crossing nrc2 or nrc2-2 to nrc3. F1s were then selfed and F2 populations were genotyped to screen for homozygous mutations in both *Nrc2* ang *Nrc3* genes.

**Supplemental Figure S3.**
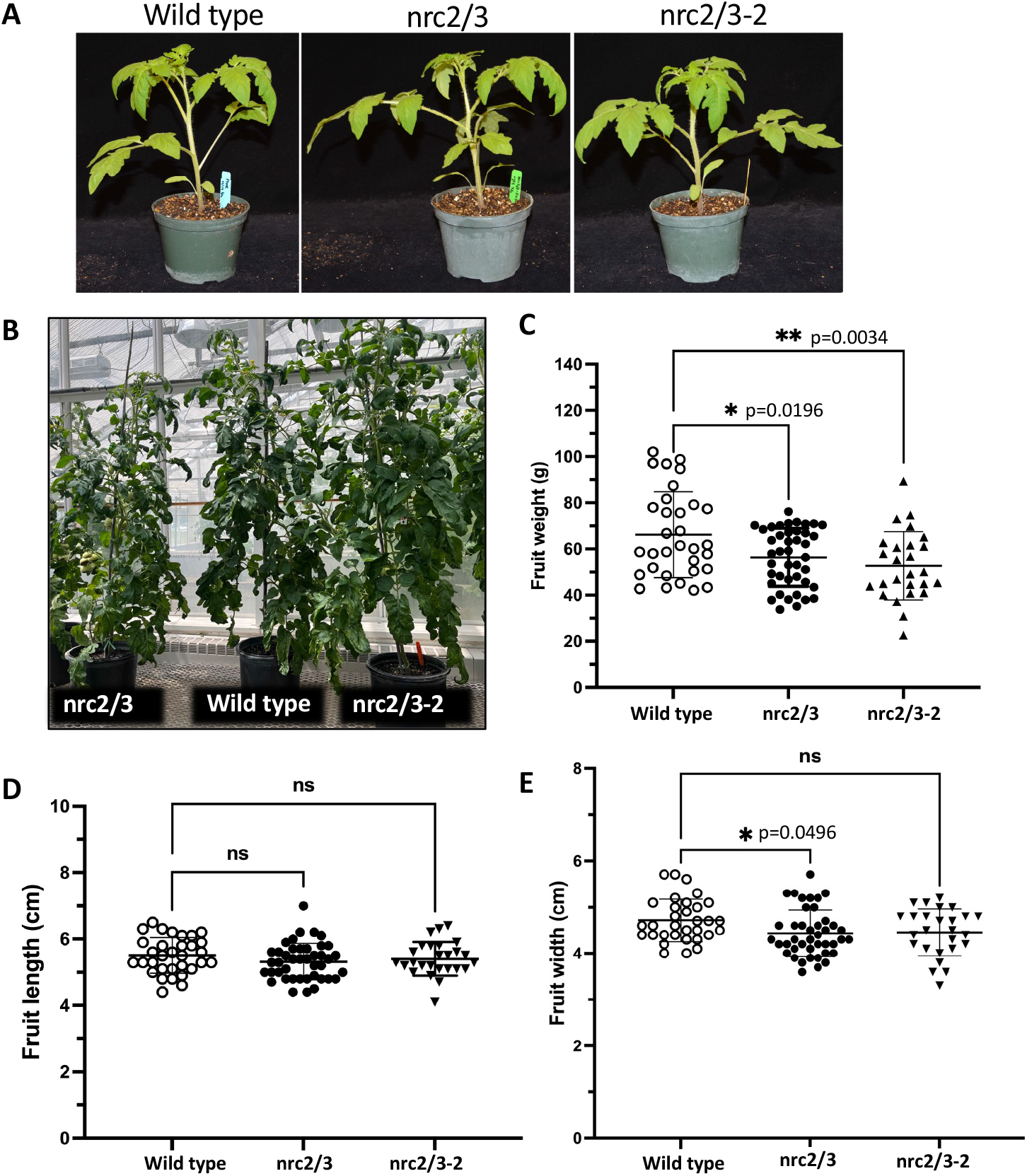
Growth, development, and morphology of the *nrc2/3* mutants were indistinguishable from wild-type RG-PtoR plants. A-B. Photographs of four-week-old or eight-week-old plants of wild-type RG-PtoR and two independent nrc2/3 mutant lines grown in the greenhouse. **C**-**E.** Average fruit weight, length and width were indistinguishable between the two nrc2/3 mutant lines and the wild-type RG-PtoR plants. Bars show means ± SD. Statistical significance was determined by pairwise t test (* p<0.05 and **p<0.01; ns, no significance).

**Supplemental Figure S4.**
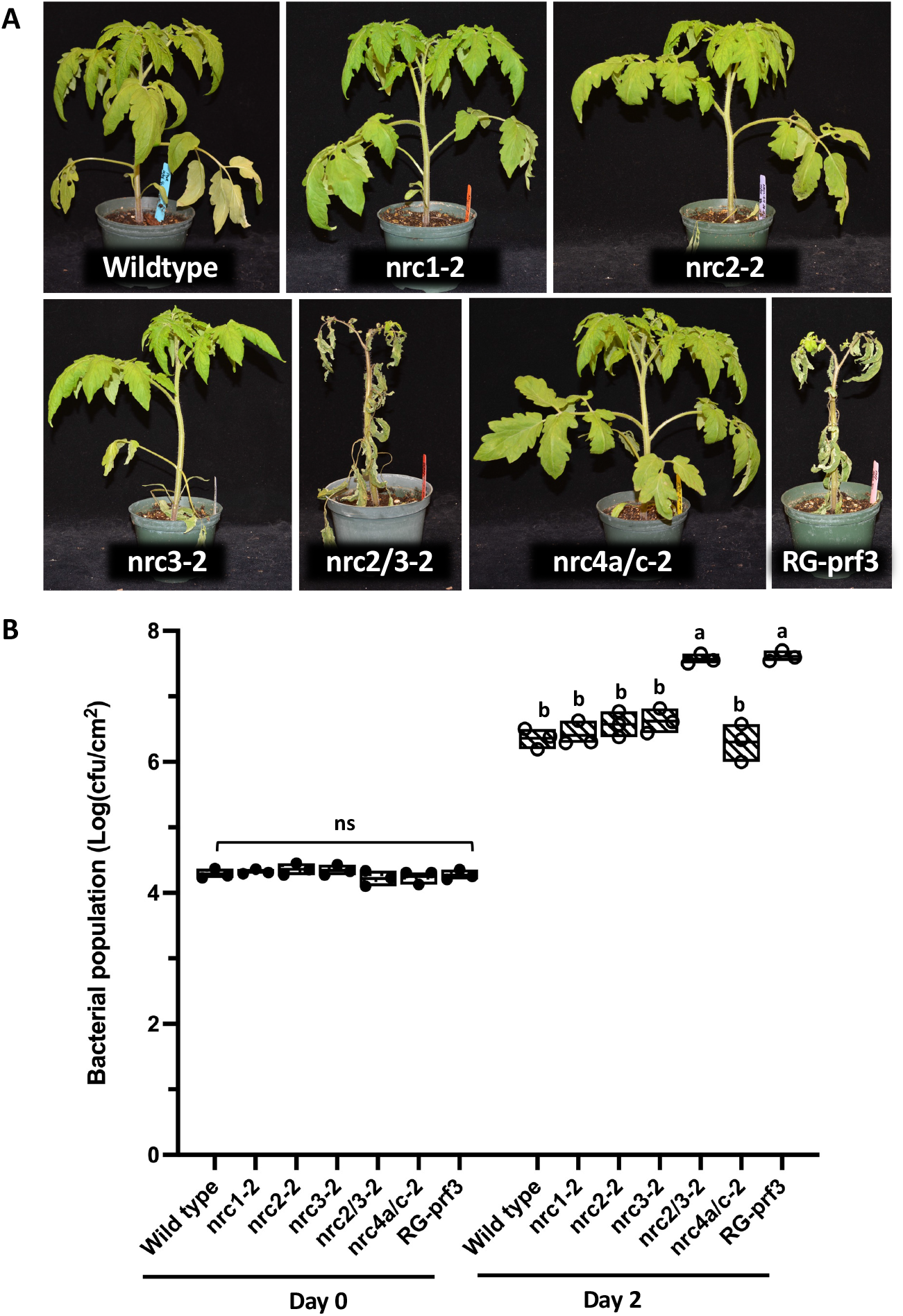
Investigation of AvrPto- and AvrPtoB-triggered immunity in the nrc mutants. **A**-**B**, Four-week-old plants of wild-type RG-PtoR, nrc1-2, nrc2-2, nrc3-2, nrc2/3-2, nrc4a/c-2 mutants, and RG-prf3 (a *Prf* mutant) plants were vacuum-infiltrated with 1 x 10^6^ cfu/mL DC3000. **A**, Photographs of disease symptoms were taken 3 days after inoculation. **B**, Bacterial populations were measured 3 hours (Day 0) and two days (Day 2) after infiltration. Bars show means ± standard deviation (SD). Different letters indicate significant differences based on a one-way ANOVA followed by Student’s *t* test (p < 0.05). ns, no significant difference. Three plants for each genotype were tested per experiment. The experiment was performed twice with similar results.

**Supplemental Figure S5.**
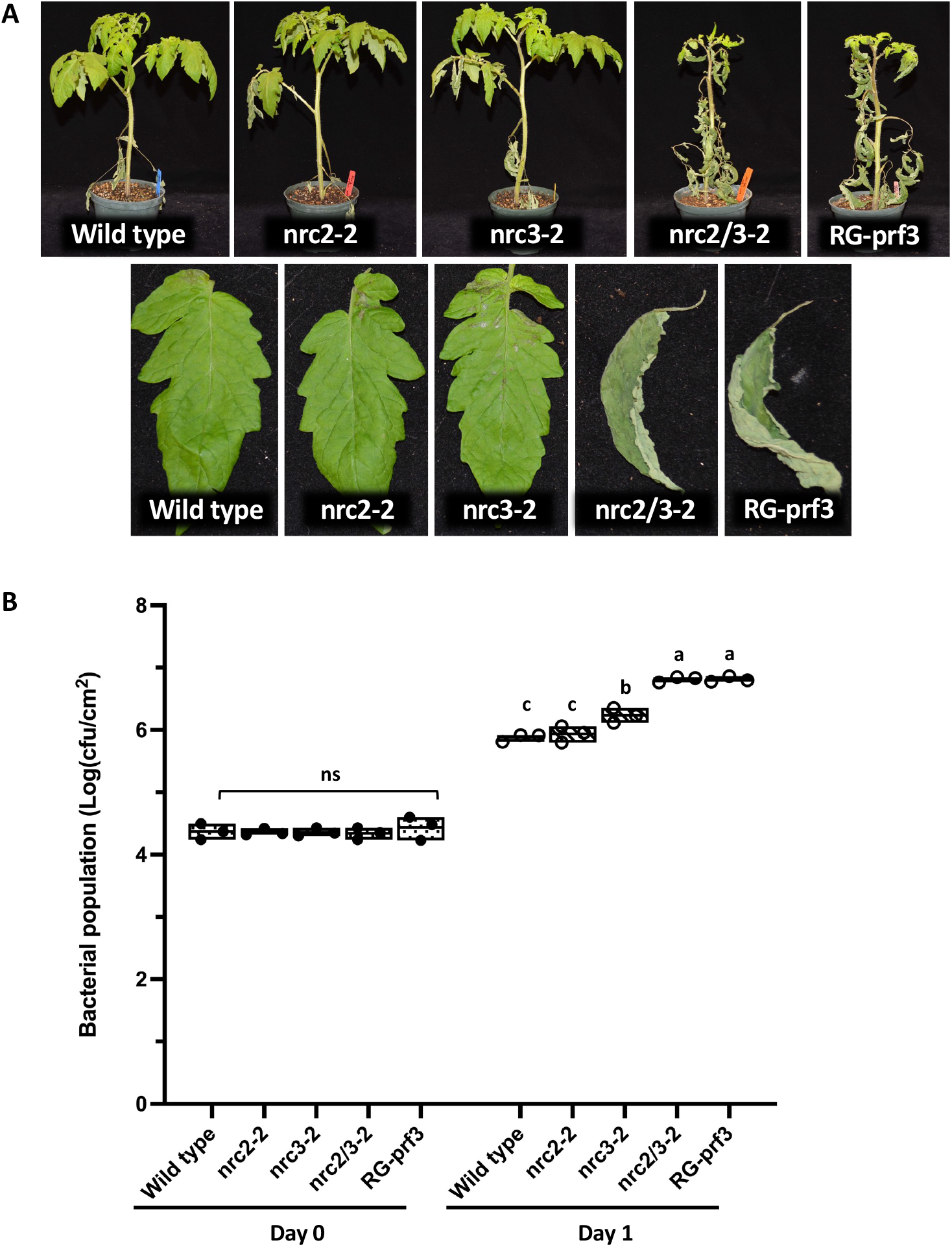
Investigation of AvrPto-triggered immunity in the nrc mutants. **A**-**B**, Four-week-old plants of wild-type RG-PtoR, nrc2-2, nrc3-2, nrc2/3-2, and RG-prf3 plants were vacuum-infiltrated with 1 x 10^6^ cfu/mL DC3000Δ*avrPtoB*. **A**, Photographs of disease symptoms of the whole plants or leaflets from the 4^th^ leaf were taken at 4 days after inoculation. **B**, Bacterial populations were measured at 3 hours (Day 0) and one day (Day 1) after infiltration. Bars show means ± SD. Different letters indicate significant differences based on a one-way ANOVA followed by Student’s *t* test (p < 0.05). ns, no significant difference. Three plants for each genotype were tested per experiment. The experiment was performed twice with similar results.

**Supplemental Figure S6.**
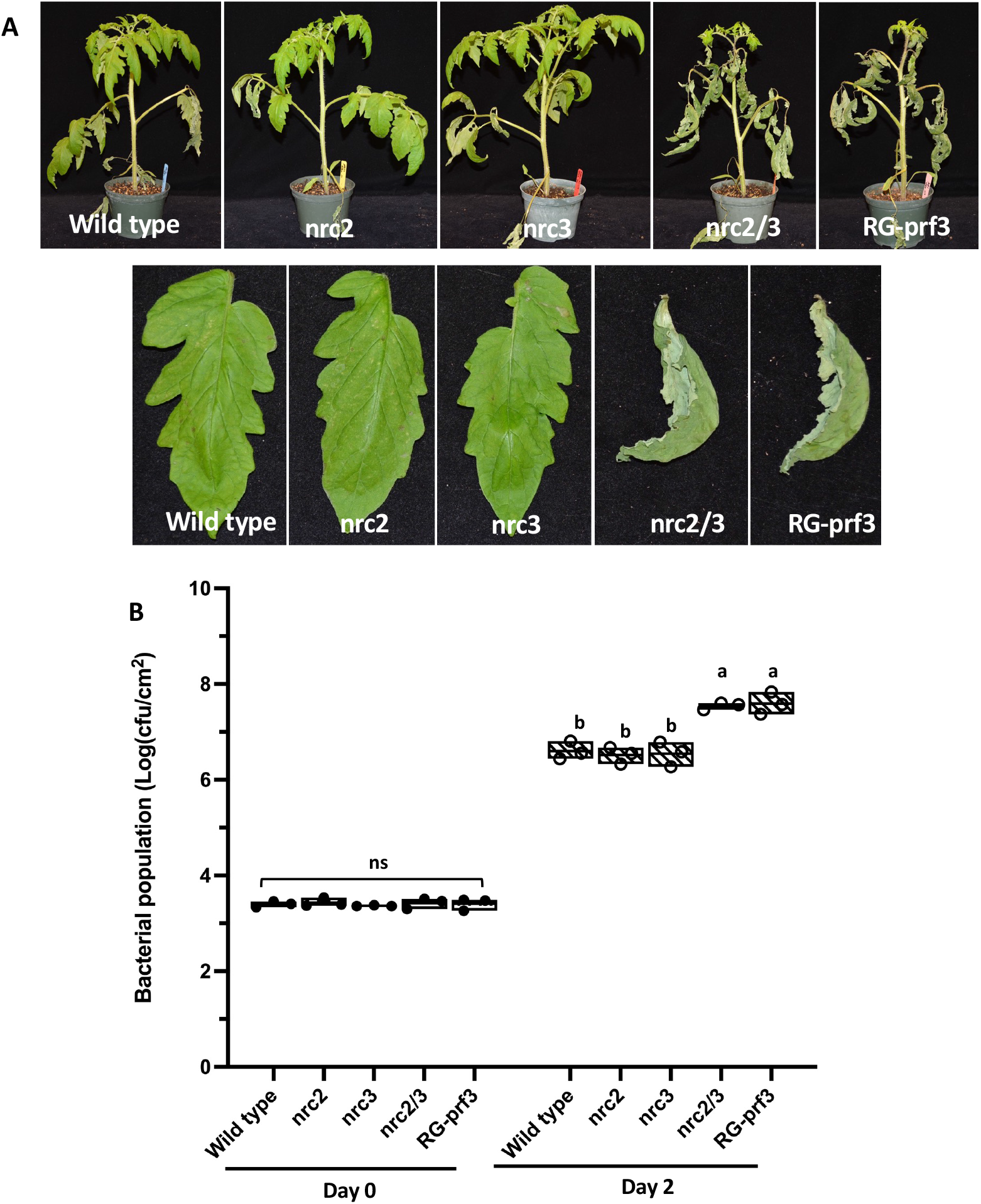
Investigation of AvrPto-triggered immunity in the nrc mutants. **A**-**B**, Four-week-old nrc2, nrc3, nrc2/3 mutants and wild-type RG-PtoR plants were vacuum-infiltrated with 1 x 10^5^ cfu/mL DC3000Δ*avrPtoB*. **A**, Photographs of disease symptoms of the whole plants or leaflets from the 4^th^ leaf were taken at 4 days after inoculation. **B**, Bacterial populations were measured at 3 hours (Day 0) and two days (Day 2) after infiltration. Bars show means ± standard deviation (SD). Different letters indicate significant differences based on a one-way ANOVA followed by Student’s *t* test (p < 0.05). ns, no significant difference. Three plants for each genotype were tested per experiment. The experiment was performed twice with similar results.

**Supplemental Figure S7.**
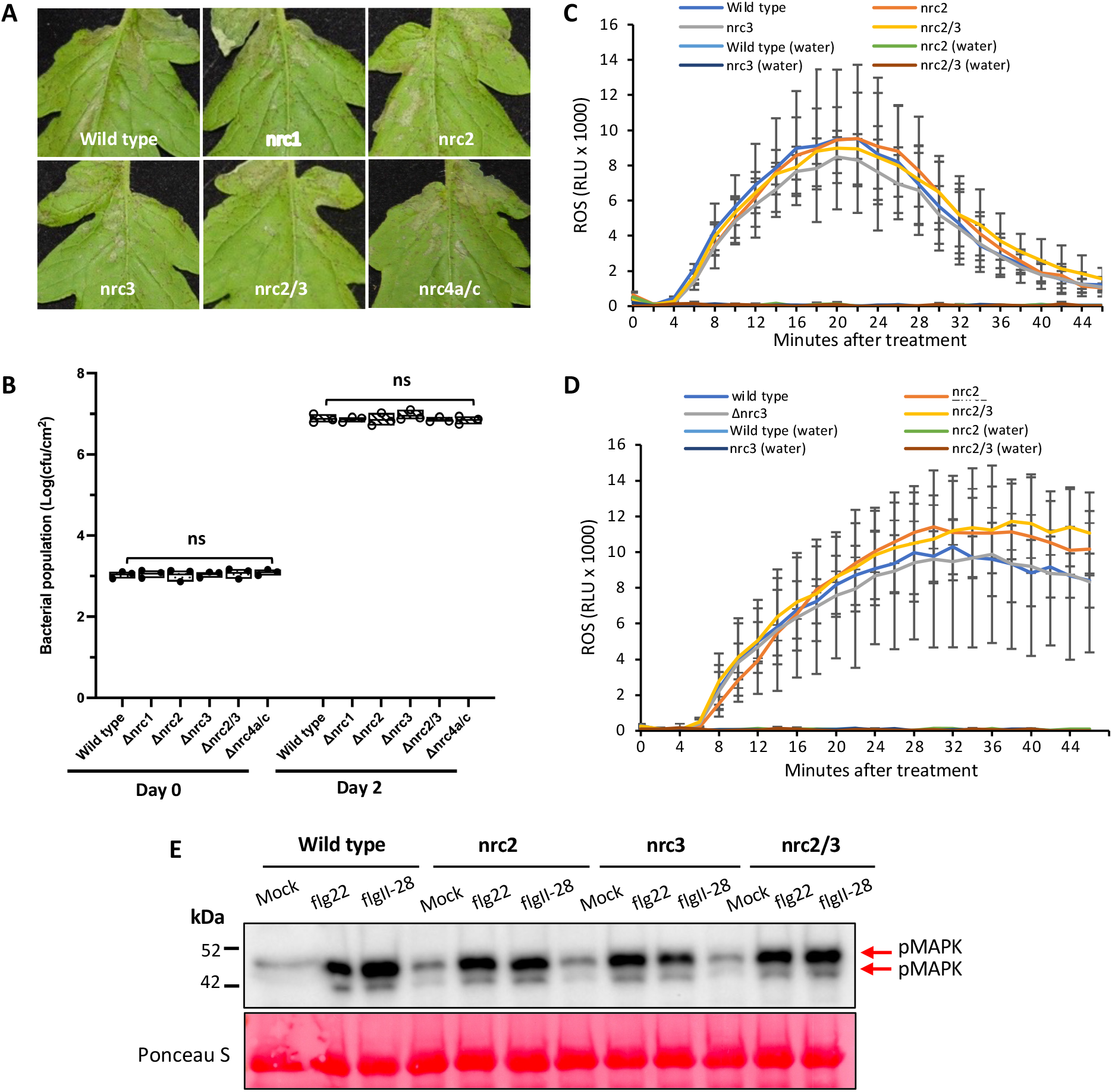
Investigation of flagellin-mediated pattern-triggered immunity in the nrc mutants. **A**-**B**, Four-week-old plants of wild-type Rio Grande (RG)-PtoR, nrc1, nrc2, nrc3, nrc2/3, and nrc4a/c mutants were vacuum-infiltrated with 5 x 10^4^ cfu/mL DC3000Δ*avrPto*Δ*avrPtoB*. **A**, Photographs of disease symptoms were taken 4 days after inoculation. **B**, Bacterial populations were measured 3 hours (Day 0) and two days (Day 2) after infiltration. Bars show means ± standard deviation (SD). Different letters indicate significant differences based on a one-way ANOVA followed by Student’s *t* test (p < 0.05). ns, no significant difference. Three plants for each genotype were tested per experiment. **C**-**D**, Leaf discs from nrc2, nrc3, nrc2/3, or wild-type RG-PtoR plants were treated with 50 nM flg22 (**C**), 50 nM flgII-28 (**D**), or water only. Relative light units (RLU) were measured over 45 minutes. One-way ANOVA followed by Student’s *t* test (p < 0.05) was performed at 24 min (peak readout) and 45 min after treatment. Bars represent means ± SD. No significant difference was observed between nrc2, nrc3, nrc2/3, and wild-type plants with either treatment. **E**, Leaf discs from wild-type RG-PtoR plants, nrc2, nrc3, or nrc2/3 mutants were treated with 20 nM flg22, 25 nM flgII-28, or water (mock) for 10 min. Proteins were extracted from a pool of leaf discs from plants and subjected to immunoblotting using an *anti*-pMAPK antibody that detects phosphorylated MAPKs (red arrows). Ponceau staining shows equal loading of proteins. These experiments was performed twice with similar results.

**Supplemental Figure S8.**
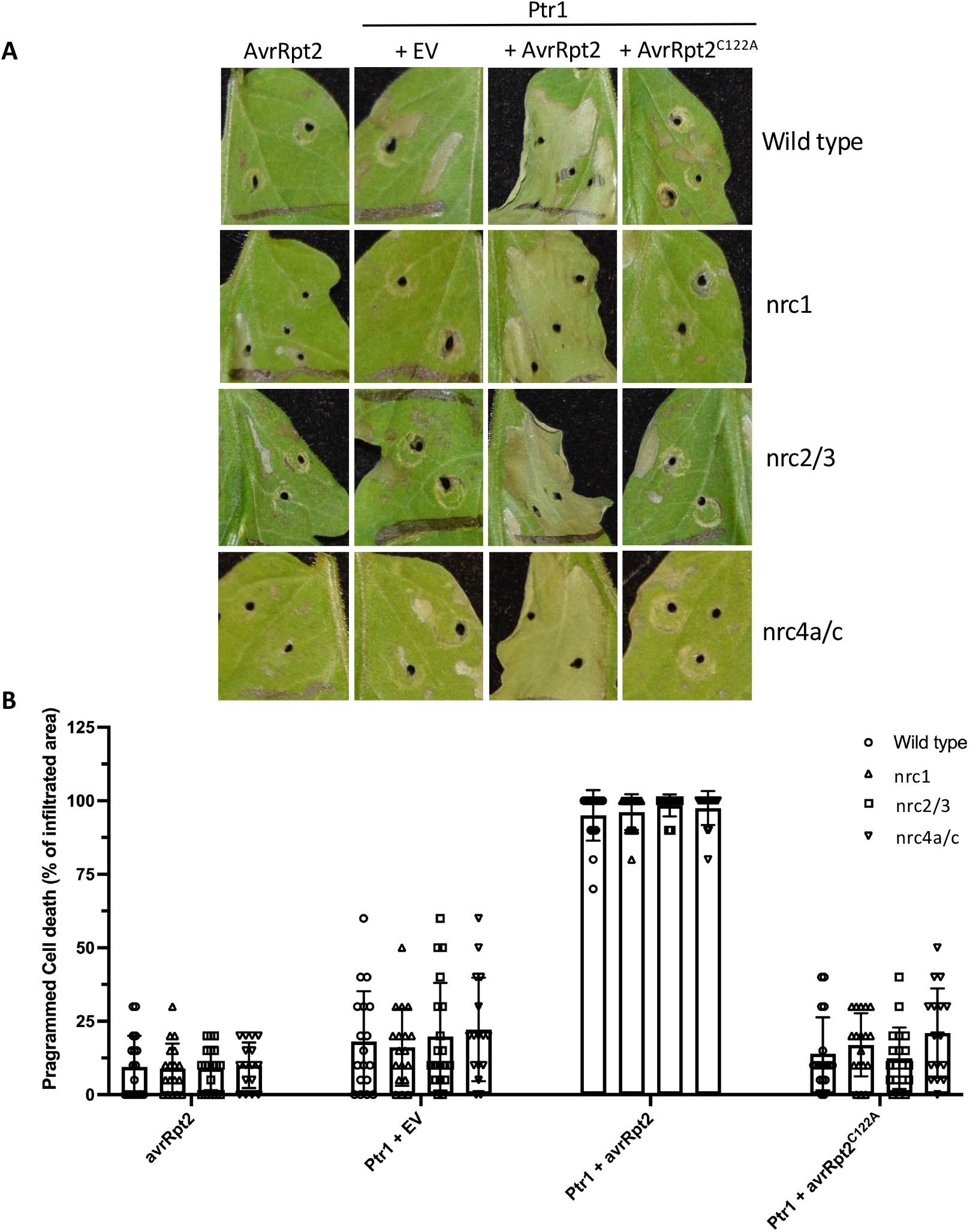
Nrc2 and Nrc3 are not required for Ptr1-mediated programmed cell death in tomato. **A**-**B**, Leaves of five-week-old plants of wild-type RG-PtoR, nrc1, nrc2/3, and nrc4 mutants were syringe infiltrated with *Agrobacterium tumefaciens* strains carrying avrRpt2 alone, or Ptr1 with empty vector (EV), avrRpt2, or avrRpt2(C122A) (OD_600_ = 0.1). **A**, Photographs were taken 4 days after infiltration. **B**, Average programmed cell death (PCD) scores. Results are based on 18 technical replicates from three biological experiments. No significant differences were observed between the wild-type plants and nrc mutants based on a one-way ANOVA followed by Student’s *t* test (p < 0.05).

**Supplemental Figure S9.**
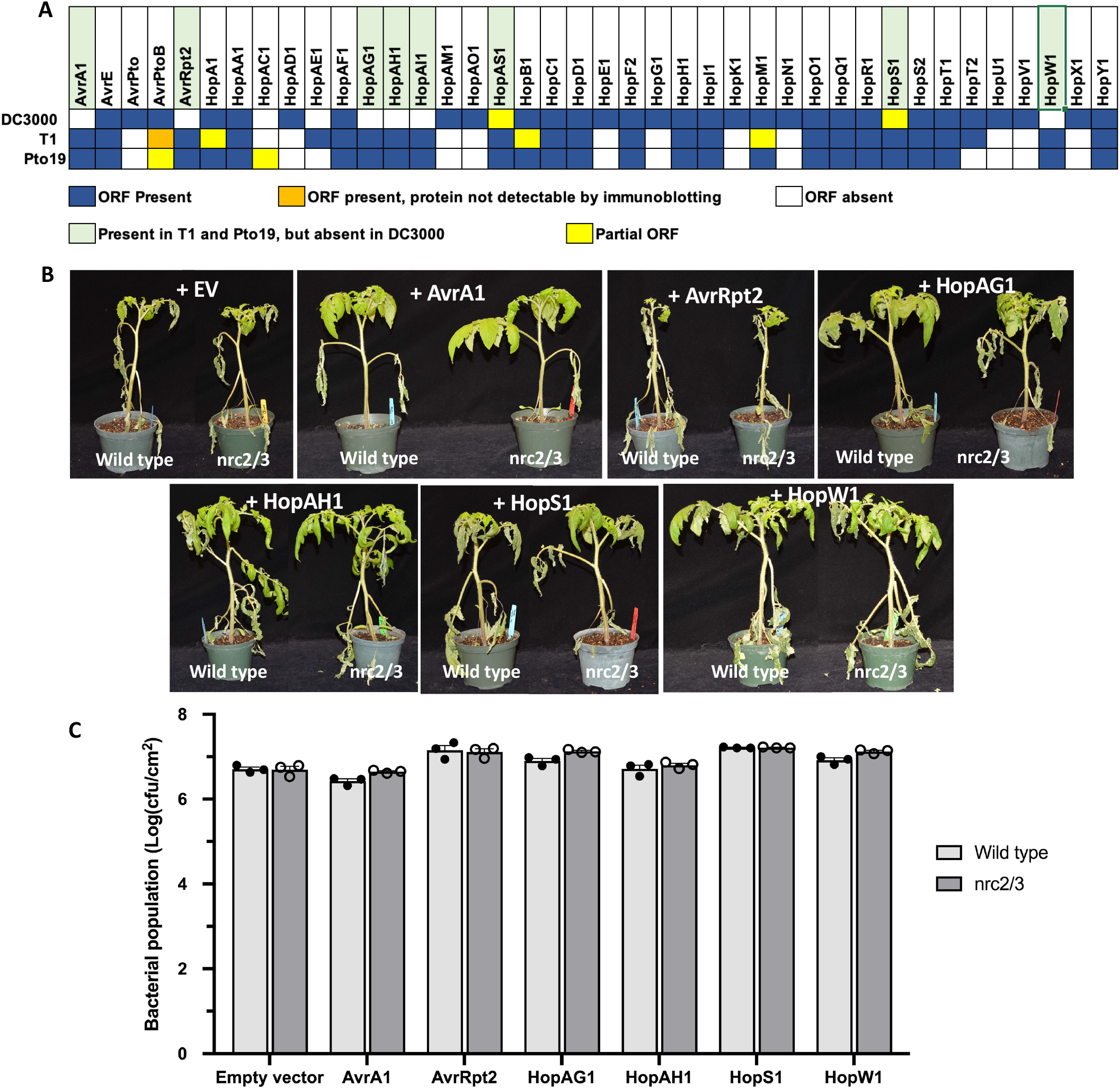
Nrc2/3 do not appear to mediate recognition of type III effectors that are present in Pto19 and absent from DC3000. **A**, Summary of the type III effectors present in the *P. syringae* strains DC3000, T1, and Pto19. A blue box indicates presence of a full length open-reading frame (ORF). An orange box indicates the gene is present but protein is not detectable by immunoblotting. A yellow box indicates the gene has an ORF truncated by an insertion sequence element or the presence of a premature stop codon. A white box indicates the gene is not present in that strain. The light green shading indicates the effectors that are present in T1 and Pto19 but absent in DC3000. **B**-**C**, Four-week-old wild-type RG-PtoR and nrc2/3 mutants were vacuum-infiltrated with DC3000Δ*avrPto*Δ*avrPtoB* carrying an empty vector (EV) or the test effector at 2 x 10^5^ cfu/mL. **B**, Photographs of disease symptoms were taken 5 days after inoculation. **C**, Bacterial populations were measured two days later. Bars show means ± standard deviation (SD). No significant differences were observed between the wild-type plants and nrc2/3 mutants based on a one-way ANOVA followed by Student’s *t* test (p < 0.05). Three plants for each genotype were tested per experiment. The experiment was performed twice with similar results.

**Supplementary Table S1.** RNA-seq data of the expression pattern of *Nrc* genes in tomato against *Pseudomonas syringae* pv. *tomato*\*.

| Gene | Solyc ID | PTI |  | ratio | p value<br>(FDR) | NTI |  | ratio | p value<br>(FDR) |
| --- | --- | --- | --- | --- | --- | --- | --- | --- | --- |
|  |  | flgII-28<br>6 h | mock<br>6 h |  |  | RG-PtoR<br>DC3000 6 h | RG-prf3<br>DC3000 6 h |  |  |
| *Nrc1* | Solyc01g090430 | 47.65 | 14.55 | 3.28 | 2.31E-11 | 62.37 | 23.99 | 2.6 | 7.39E-13 |
| *Nrc2* | Solyc10g047320 | 83.23 | 31.46 | 2.65 | 8.96E-07 | 30.91 | 24.07 | 1.28 | 0.1477 |
| *Nrc3* | Solyc05g009630 | 160.86 | 86.34 | 1.86 | 0.0016 | 90.04 | 118.39 | 0.77 | 0.4008 |
| *Nrc4a* | Solyc04g007070 | 99.07 | 36.72 | 2.7 | 9.07E-05 | 56.83 | 28.91 | 1.97 | 1.69E-07 |
| *Nrc4c* | Solyc04g007030 | 10.15 | 3.41 | 2.97 | 4.68E-08 | 5.41 | 3.15 | 1.72 | 0.0506 |

\*Numbers under PTI and NTI are RPKMs (reads per kilobase of transcript per million mapped reads). FDR, false discovery rate. Data are available on the Tomato Functional Genomics Database (TGFD; http://ted.bti.cornell.edu/cgi-bin/TFGD/digital/home.cgi). Data (RPKM from RNA-Seq) for pattern-triggered immunity (PTI) are from Rosli et al., (2013) and for NLR-triggered immunity (NTI) are from Pombo et al., (2014).

**Supplemental Table S2.**
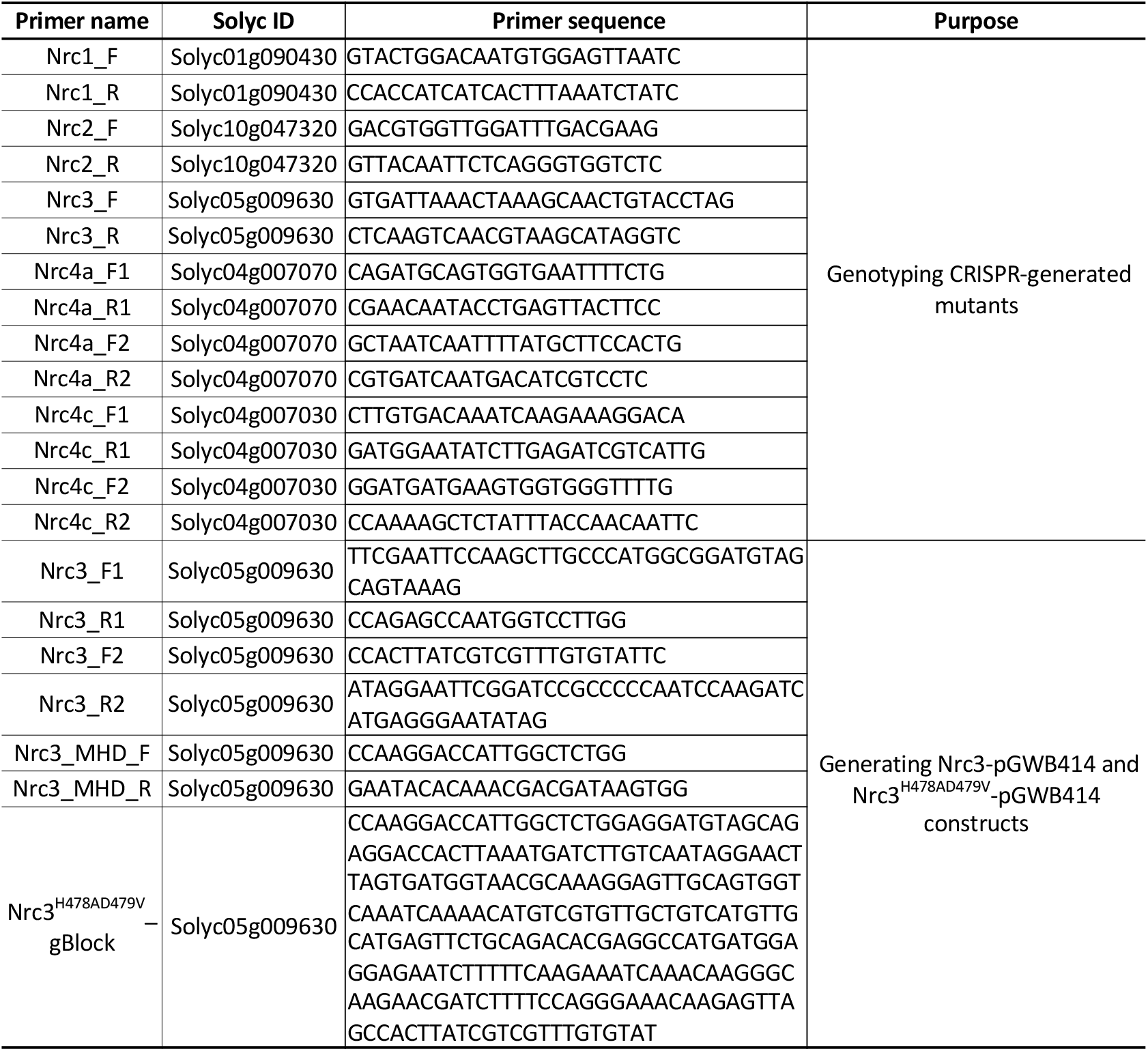
Primers used for genotyping CRISPR-generated mutants.

**Supplemental Table S3.** Summary of *Pseudomonas syringae* pv. *tomato* (*Pst*) strains.

| ***Pst* strain** | **Selective marker** | **Reference** |
| --- | --- | --- |
| DC3000 | Rif | Buell et al., 2003 |
| DC3000Δ*avrPto* | Rif/Spec | Ronald et al. 1992 |
| DC3000Δ*avrPtoB* | Rif/Kana | Lin and Martin, 2005 |
| DC3000Δ*avrPto*Δ*avrPtoB* | Rif | Lin and Martin, 2005 |
| DC3000Δ*avrPto*Δ*avrPtoB*Δ*fliC* | Rif | Alan Collmer |
| T1Δ*avrPtoB*Δ*fliC* | Rif | Almeida et al., 2009 |
| Pto19Δ*avrPtoB* | Spec | Kunkeaw et al., 2010 |

**Supplemental Table S4.** Plasmids and *Agrobacterium* strains.

| **Autoactive protein** | **Plasmid** | **Selective marker** | **Agrobacterium strain** | **Reference** |
| --- | --- | --- | --- | --- |
| AvrPtoB<sub>1-387</sub> | pGWB414 | Spec | 1D1249 | This study |
| Pto<sup>Y207D</sup> | pBTEX | Kan | 1D1249 | Du et al., 2012 |
| Prf<sup>D1416V</sup> | pBTEX | Kan | 1D1249 | Du et al., 2012 |
| M3Ka<sup>KD</sup> | pER8 | Spec | GV2260 | Oh et al., 2010 |
| Mkk2<sup>DD</sup> | pER8 | Spec | GV2260 | Pedley and Martin, 2004 |
| Nrc3<sup>H478AD479V</sup> | pGWB414 | Spec | GV2260 | This study |

## Notes

### Competing Interest Statement

The authors have declared no competing interest.

